# IL-17A Alters Human Cortical Development in a 3D Ex Vivo Model of Maternal Immune Activation

**DOI:** 10.1101/2024.10.30.621011

**Authors:** Muhammad Z. K. Assir, Mario Yanakiev, Do Hyeon Gim, Sara S. M. Valkila, Paola Muscolino, Liu Peng, Paul A. Fowler, Daniel A. Berg, Eunchai Kang

## Abstract

Human brain development depends on the coordinated interaction of diverse cell types and extracellular matrix (ECM) components, which are essential for proper neurogenesis and cortical organization. Epidemiological and animal studies have demonstrated that maternal immune activation (MIA) disrupts brain development, leading to impaired neurogenesis and increased risk of neurodevelopmental disorders (NDDs), including autism spectrum disorder (ASD) and schizophrenia. However, the cellular and molecular mechanisms by which MIA impacts human cortical development remain poorly understood. Here we introduce a 3D ex vivo culture system, termed ‘cerebroids,’ derived from dorsolateral prefrontal cortex of human fetal brain tissue, which faithfully preserves key developmental processes, along with critical cellular diversity and structural integrity of the developing human cortex. Using this platform, we show that IL-17A, a cytokine strongly implicated in NDDs, induces premature cortical folding, increases cortical thickness, and accelerates neurogenesis and neuronal maturation. Transcriptomic and proteomic analyses reveal significant dysregulation of ECM-related pathways, including the upregulation of proteoglycans such as brevican and versican. Notably, treatment with the anti-inflammatory agent parthenolide, an inhibitor of NF-κB and HDAC1 pathways, reverses IL-17A-induced cortical abnormalities, restoring normal cortical thickness, folding, and neurogenesis. These findings provide valuable insights into how IL-17A disrupts human cortical development during MIA, advancing our understanding of NDD-associated structural cortical alterations.

## Main Text

A growing body of evidence from epidemiological studies has demonstrated that maternal immune activation (MIA) significantly increases the risk of neurodevelopmental disorders (NDDs), including autism spectrum disorder (ASD) and schizophrenia, in offspring (*1-5*). Animal models of MIA have provided insights into its role in the pathogenesis of NDDs, revealing ASD-related behavioral deficits and structural brain abnormalities (*6, 7*), such as increased cortical thickness, elevated neuronal numbers (*8*), and perturbed cortical layer organization (*7*). Despite these findings, the precise cellular and molecular mechanisms by which the inflammatory environment in utero alters the developmental trajectory of the human cortex, leading to such structural and functional abnormalities, remain poorly understood.

Human pluripotent stem cell (PSC)-derived brain organoids offer a valuable platform for investigating human-specific aspects of developmental disturbances caused by genetic and environmental insults (*9*). However, the absence of immune and vascular components in these organoid models restricts their use, particularly in modeling MIA (*10*). Ex vivo models using primary human tissue provide a more accurate representation of brain development, but conventional slice cultures are limited by low throughput and scalability issues. To address this critical gap, we have developed a high-throughput ex vivo three-dimensional (3D) culture system, called ‘cerebroids’, using human fetal brain tissue.

### Establishment and validation of a 3D ex vivo culture of human fetal brain

To establish the 3D ex vivo culture system of the human fetal brain, we dissected the dorsolateral prefrontal cortex (DLPFC) from healthy fetal brain tissues at 11-14 gestational weeks (GW), a critical period of active neurogenesis in the developing cortex (11) and when MIA is most strongly associated with an increased risk of NDDs **(fig. S1, A and B, table S1)** (*3*). The tissue was cut into approximately 10 mm^2^ pieces, ensuring that each section retained intact architecture from the ventricular surface to the pial surface. These pieces were cultured in a free-floating manner on an orbital shaker **(Fig. 1A, and fig. S1, C and D)**. This culture method preserves the structural integrity of the tissue and provides versatility for multiple downstream applications, including immunofluorescence staining on cryosections and comprehensive multiomics analyses **(Fig. 1A, and fig. S1E)**.

**Fig 1.**
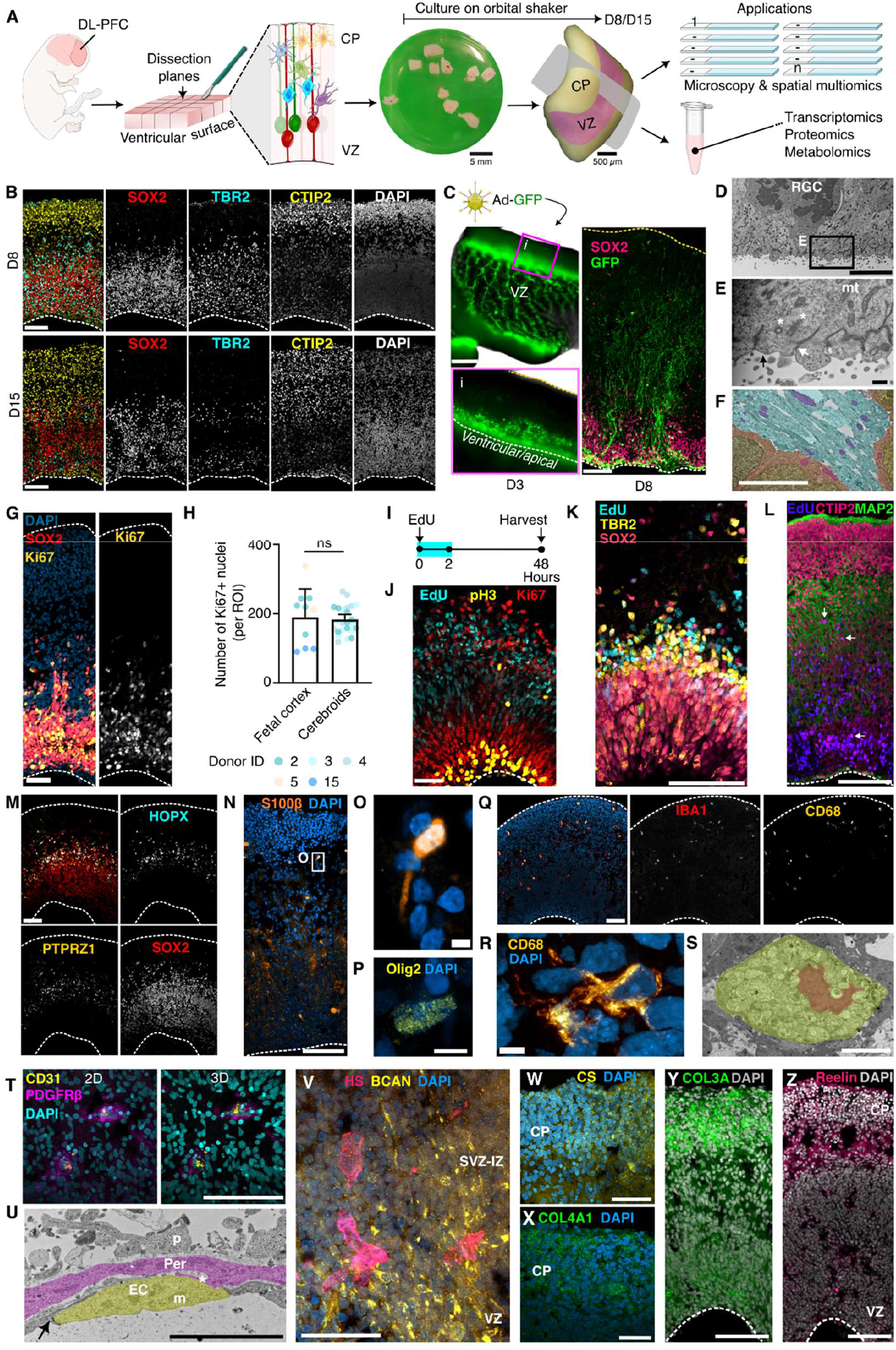
Establishment of a 3D culture method for human fetal brain tissue. **(A)** Schematic of the experimental paradigm. The dorsolateral prefrontal cortex (DL-PFC) was dissected, cultured, and processed for analysis. CP, cortical plate; VZ, ventricular zone. **(B)** Representative immunofluorescence images of cerebroids cultured for 8 and 15 days, showing the layered organization of developing brain tissue. Dotted lines indicate the ventricular surface. **(C)** Left: Wholemount live image of Adenovirus-GFP-infected cerebroids after 3 days in culture. Right: Immunofluorescence image of a section from day 8 cerebroids showing GFP^+^ neural stem cells (NSCs). **(D** to **F)** Transmission electron microscopy (TEM) images of day 2 cerebroids illustrating NSCs lining the ventricular surface (D), higher-magnification view of the apical surface showing mitochondria (mt), centrioles (*), tight junction (white arrow), and secretory vesicles (black arrow) (E). Pseudo-colored TEM image illustrating radial glia cell bodies (brown) and processes (cyan), enriched with mitochondria (purple) (F). **(G)** Immunofluorescence image of proliferating (Ki67^+^) NSCs in cerebroids. **(H)** Quantification of the number of Ki67^+^ cells in fetal cortex versus day 8 cerebroids. Data represented as mean ± SEM. ns: not significant (p ≥ 0.05). **(I** to **L)** EdU pulse-chase experiment on cerebroids. Schematic of the EdU pulse-chase experimental design (I), maximum intensity projection image showing mitotic (pH3^+^), proliferating (Ki67^+^), and EdU^+^ cells (J), EdU^+^TBR2^+^ intermediate progenitors (K), and EdU^+^ CTIP2^+^ and Map2^+^ neurons (arrows) in day 2 cerebroids (L). **(M)** Representative image of HOPX and PTPRZ1 expression in outer radial glia (oRGs) cells in day 8 cerebroid. **(N** to **R)** Immunofluorescence images of different cell types in cerebroids: S100β^+^ astroglia progenitors (N and O), Olig2^+^ oligodendrocyte precursor cells (OPCs) (P), and IBA1^+^ and CD68+ microglia (Q and R). **(S)** TEM image of a microglial cell, with pseudo-coloring of the nucleus (brown) and cell body (yellow). **(T)** Immunofluorescence image of PDGFRβ^+^ pericytes and CD31^+^ endothelial cells in cultured tissue. **(U)** TEM image of a blood vessel in day 2 cerebroids, showing endothelial cells (EC), tight junction (arrow), pericytes (Per), pericyte peg-sockets (*), radial glia processes (p), and mitochondria (m) (also see fig. S3, D–F for higher magnification images of the same). **(V to Z)** Fluorescence images show key extracellular matrix (ECM) components. Abbreviations: HS, heparan sulfate; BCAN, brevican; CS, chondroitin sulfate; SVZ-IZ, subventricular and intermediate zones. Scale bar: 500 μm (C, left), 100 μm [(B), (C, right), (G), (J to N), (Q), (T), (V to Z)], 5 μm [(D), (F), (O), (P), (R), (S), (U)], and 500 nm (E).

The cytoarchitecture and apical-basal polarity of cells in the developing brain, particularly neural stem cells (NSCs) are critical for the proper developmental trajectory of the cortex (*11*). Cerebroids cultured for 8 and 15 days retained the organized layers of SOX2^+^ NSCs, TBR2^+^ intermediate progenitor cells (IPCs) and CTIP2^+^ neurons **(Fig. 1B)**. The proper apical-basal polarity of NSCs and the integrity of their processes were confirmed by adenovirus-mediated GFP labelling of NSCs **(Fig. 1C)**, Nestin and GFAP immunostaining **(fig. S2, A and B)**, as well as scanning electron microscopy (SEM) imaging **(fig. S2, C to E)**. Notably, unlike cortical organoids derived from human induced pluripotent stem cells (iPSCs), the expression pattern of GFAP in NSCs within cerebroids was comparable to that of acutely fixed fetal tissue. The integrity of apical junctions in NSCs was verified by transmission electron microscopy (TEM) **(Fig. 1, D to F)**. Optimal culture conditions that ensure proper proliferation rate of NSCs are crucial for maintaining the polarity between the ventricular and cortical surfaces (*12*). In cerebroids, NSCs exhibited a proliferative capacity comparable to that of acutely fixed fetal cortex, as demonstrated by Ki67 staining **(Fig. 1, G and H)**. Furthermore, the capacity of NSCs to generate IPCs and neurons was confirmed through EdU pulse-chase experiments **(Fig. 1, I to L)**. Mitotic nuclei of NSCs (pH3+) were also observed near the ventricular surface, mirroring the behavior of NSCs undergoing interkinetic nuclear migration during cell division in the developing cortex. **(Fig. 1J)**. Next, we verified the presence of different cell types in cerebroids including HOPX^+^ or PTPRZ1^+^ outer radial glial cells, CTIP2^+^ or SATB2^+^ neurons, S100β^+^ or Olig2^+^ macroglia precursor cells, and IBA1^+^/CD68^+^ microglia **(Fig. 1M and fig. S3, A to C)**. We also confirmed the presence of vascular structures containing endothelial cells and pericytes **(Fig. 1, T to U, fig. S3, D to H)**.

The extracellular matrix (ECM) creates a dynamic environment that facilitates interactions among various cell types by modulating the diffusion and distribution of signaling molecules. It is also involved in the regulation of central developmental processes, such as cell proliferation, differentiation, and migration (*12*). We confirmed that cerebroids maintain key components of the ECM in the developing cortex, such as heparan sulfate, brevican, chondroitin sulfate, collagens, and ECM-associated protein Reelin **(Fig. 1, V to Z)**. Importantly, cerebroids, in contrast to iPSCs-derived cortical organoids, showed expression patterns of heparan sulfate and chondroitin sulfate that were comparable to acutely fixed fetal tissue **(fig. S4, A and B)**.

Together, these findings demonstrate that cerebroids offer a robust platform that preserves key cellular architecture, along with immune, vascular, and essential ECM components, enabling the study of both healthy neurodevelopment and disruptions caused by environmental factors.

### Abnormal cortical development mediated by IL-17A

Emerging evidence links IL-17A to NDDs. Elevated levels of IL-17A are correlated with ASD severity in humans (*13-15*) and genetic variations in the IL17A gene have been identified in ASD cohorts (*16*). In animal models, IL-17A mediates MIA-induced autism-like behaviors and cortical abnormalities, which can be prevented by blocking IL-17A activity (*7, 17, 18*). To further explore the impact of IL-17A during MIA in cortical development (for additional rationale on using IL-17A to model MIA, see the Supplementary Text), we utilized cerebroids from GW 11-14 fetal brain tissues. Transcriptomic analysis of RNA-sequencing data from microglia/brain associated macrophages (BAM) isolated from cerebroids revealed that IL-17A treatment in culture activated various inflammation-related pathways, such as TLR, TNF, and NF-κB signaling **(Fig. 2, A and B)**. All donors used in this study exhibited comparable baseline inflammatory states, as determined by transcriptomic analysis of inflammatory markers in placental tissue **(fig. S5)**. IL-17A-treated cerebroids showed premature induction of cortical folding at 8 days ex vivo and the folding phenotype became more prominent at 15 days ex vivo **(Fig. 2C and fig. S6, A and B)**. SEM imaging further confirmed alterations in the cortical surface structure of IL-17A-treated cerebroids **(Fig. 2D)**. To measure the degree of folding, we calculated the gyrification index (GI), the ratio of the fiducial surface length to the convex hull length, and it was found that IL-17A-treated cerebroids exhibited increased GI **(Fig. 2, E and F, and fig. S6C)**. Moreover, IL-17A-treated cerebroids exhibited a significant increase in overall thickness, which was observed across all three cortical layers **(Fig. 2, E and G, and fig. S6, D to F)**. Particularly, the IZ/SP (layer 2) showed reduction in cell density, suggesting a potential expansion of ECM in IL-17A-treated cerebroids **(Fig. 2E)**. Next, we investigated which cell types contributed to the increased thickness. The number of CTIP2^+^ deep layer neurons was significantly higher in IL-17A-treated cerebroids, while no significant difference was observed in the number of SOX2^+^ NSCs or TBR2^+^ IPCs **(fig. S6G and Fig. 2, H and I)**. Additionally, IL-17A treatment increased the number of SATB2^+^ upper layer neurons and NeuN^+^ mature neurons **(Fig. 2, J to N)**, suggesting accelerated neuronal maturation in IL-17A-treated cerebroids. The increased number of excitatory neurons induced by IL-17A is consistent with previous findings from studies modeling MIA using IL-6 in human dorsal forebrain organoids (*19*). To further confirm this, we treated neurons differentiated from iPSC-derived neural progenitor cells (NPCs) with IL-17A and performed Ca^2+^ imaging. IL-17A-treated neurons exhibited increased spontaneous neuronal activity, further supporting enhanced maturation **(fig. S7, A to C)**. Premature cortical folding, increased cortical thickness, and accelerated neuronal maturation mirror clinical observations in some individuals with ASD, including macrocephaly (*20, 21*), an increased number of prefrontal cortical neurons (*22*), and enhanced cortical folding (*23*).

**Fig 2.**
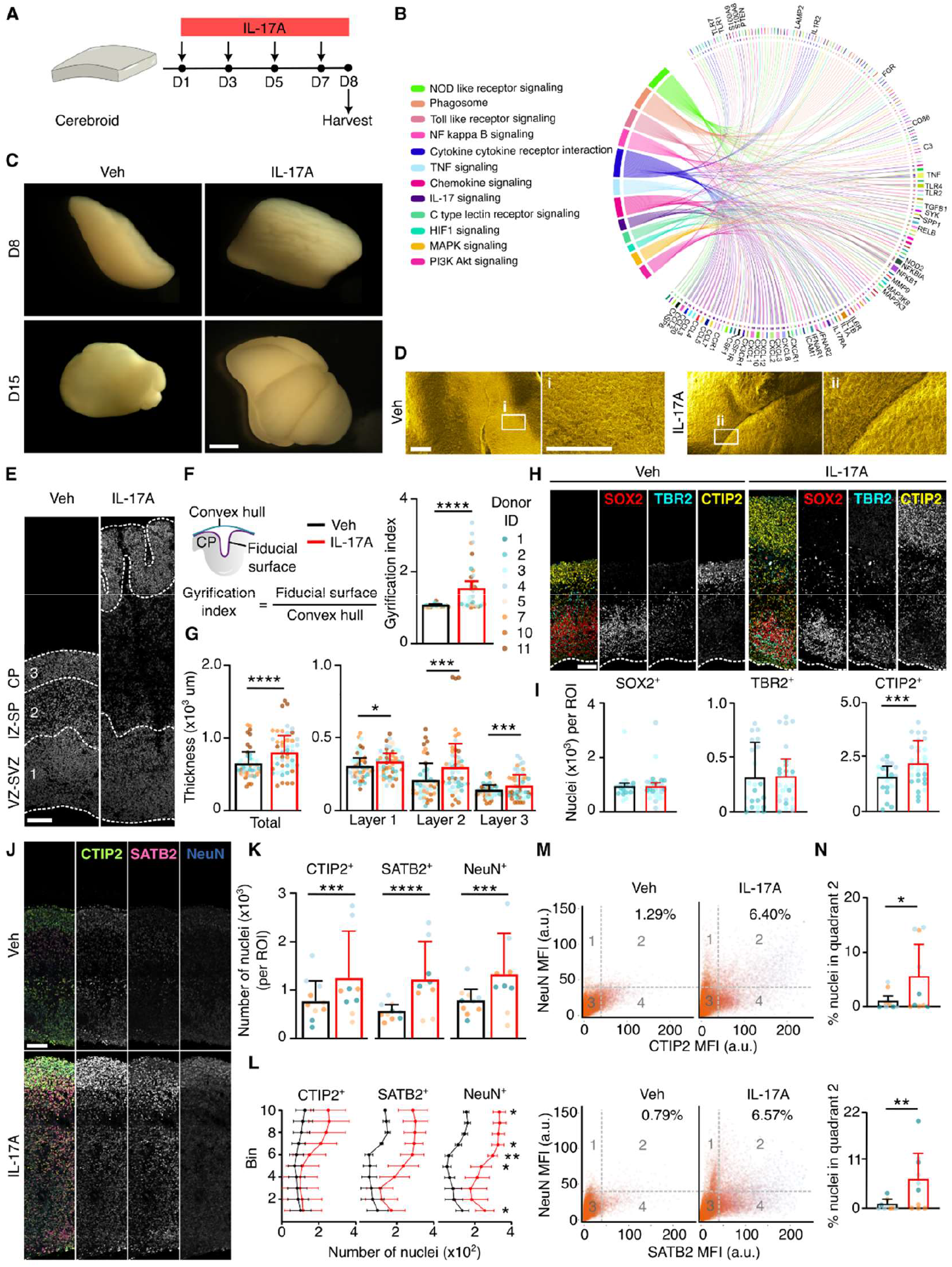
IL-17A treatment promotes cortical folding and increases neuronal numbers in cerebroid cultures. **(A)** Schematic illustration of the experimental paradigm for IL-17A treatment. **(B)** Chord plot showing upregulated genes (right) and their associated KEGG pathways (left) in microglia following IL-17A treatment. **(C)** Representative images of cerebroids cultured for 8 (D8) and 15 days (D15), demonstrating morphological changes. **(D)** Scanning electron microscopy (SEM) images of the cortical surfaces in vehicle- and IL-17A-treated cerebroids, demonstrating structural difference. **(E)** DAPI-stained cross-sections of vehicle- and IL-17A-treated cerebroids, showing alterations in cortical architecture. The layers shown are ventricular and subventricular zones (VZ-SVZ, Layer 1), intermediate zone and subplate (IZ-SP, Layer 2), and cortical plate (CP, Layer 3). **(F)** Schematic of gyrification index calculation (left) and quantification (right), demonstrating increased cortical folding in IL-17A-treated cerebroids (n = 18 cerebroids) compared to vehicle controls (n = 17 cerebroids) (data from 7 donors). **(G)** Quantification of total and layer-specific cortical thickness in vehicle-(n = 17 cerebroids) and IL-17A-treated (n = 18 cerebroids) cerebroids. (n = 37 cerebroids from 7 donors). (data from 7 donors). **(H)** Representative fluorescence images of neural stem cells (NSCs, Sox2), intermediate progenitor cells (IPCs, TBR2), and neurons (CTIP2) in vehicle- and IL-17A-treated cerebroids, demonstrating shifts in cell populations. **(I)** Quantification of NSC, IPC, and neuronal numbers from panel (H) in vehicle-(n = 9 cerebroids) and IL-17A-treated (n = 9 cerebroids) cerebroids. (data from 3 donors). **(J** and **K)** Representative confocal images (J) and quantification (K) of neurons expressing CTIP2, SATB2, and NeuN in vehicle-(n = 10) and IL-17A-treated (n = 9) cerebroids. (data from 4 donors). **(L)** Quantification of neuronal numbers across cortical layers (bins 1–10, from the ventricular to pial surface), showing spatial distribution in vehicle-(n = 9) and IL-17A-treated (n = 9) cerebroids (data from 3 donors). **(M)** Scatter plots displaying mean fluorescence intensities (MFI, arbitrary units) of NeuN, CTIP2, and SATB2 across nuclei per region of interest (ROI) in vehicle-(n = 9 cerebroids) and IL-17A-treated cerebroids (n = 10 cerebroids, from 3 donors). **(N)** Quantification of percentage of CTIP2^+^ and SATB2^+^ neuronal populations with high NeuN expression per ROI, corresponding to panel M. Data are presented as mean ± SEM (*p < 0.05, **p < 0.01, ***p < 0.001, ****p < 0.0001, two-way ANOVA for treatment-group and donor). Scale bar, 1000 μm (C) and 100 μm [(D), (E), (H), (J)].

### Enhanced neurogenesis by IL-17A

To identify the cellular mechanisms underlying abnormal cortical development, we examined the acute impact of IL-17A on NSCs. To determine whether the increased number of CTIP2^+^ neurons in IL-17A-treated cerebroids was due to elevated nascent neurogenesis, we conducted an EdU pulse-chase experiment. Following a 2-hour pulse and 48-hour chase, we observed a higher number of EdU^+^CTIP2^+^ neurons in IL-17A treated cerebroids, suggesting that IL-17A increases nascent neurogenesis **(Fig. 3, A to C)**. This phenotype was driven by an increased generation of TBR2^+^ IPCs, as we observed a higher number of EdU^+^TBR2^+^ cells in IL-17A treated cerebroids **(Fig. 3, D and E)**. The prolonged course of neurogenesis is a key factor in cortical expansion in humans (*24*), and premature differentiation of neural progenitors into neurons can deplete the progenitor pool, resulting in reduced neuronal numbers and often leading to a microcephaly phenotype (*25*). Although we detected more EdU-labelled CTIP2^+^ neurons and TBR2^+^ IPCs in IL-17A-treated cerebroids, the number of EdU^+^ SOX2^+^ NSCs remained comparable between the two groups **(Fig. 3, D and F)**, suggesting that enhanced neurogenesis was accompanied by increased proliferation and replenishment of the NSC population. Notably, while no difference in the number of Ki67^+^ proliferating NSCs was found between vehicle and IL-17A-treated groups, a greater number of EdU^+^ cells and EdU^+^Ki67^+^ cells were observed in IL-17A-treated cerebroids, indicating that a higher number of EdU^+^ cells reentered the cell cycle. **(Fig. 3, G to J)**. To further support the observation of increased neurogenesis induced by IL-17A, iPSC-derived cortical organoids were treated with IL-17A. Consistent with findings in cerebroids, the number of CTIP2^+^ neurons was increased in the IL-17A-treated organoids **(Fig. 3, K to M)**. However, we did not observe a cortical folding phenotype in IL-17A-treated organoids, suggesting that ECM components and cell types absent in organoids may be crucial in mediating this phenotype. Additionally, IL-17A-treatment of iPSC-derived NPCs resulted in accelerated neurogenesis, as shown by an increased number of MAP2^+^ cells in the IL-17A-treated group **(Fig. 3, N to P)**. These findings from our organoid and NPC experiments suggest that the increased neurogenesis induced by IL-17A is due to a direct effect on NSCs, rather than being mediated by microglia-driven immune responses. This contrasts with recent findings in mice, where transcriptomic changes in both neuronal and non-neuronal cells in response to MIA were shown to require microglia (*26*).

**Fig 3.**
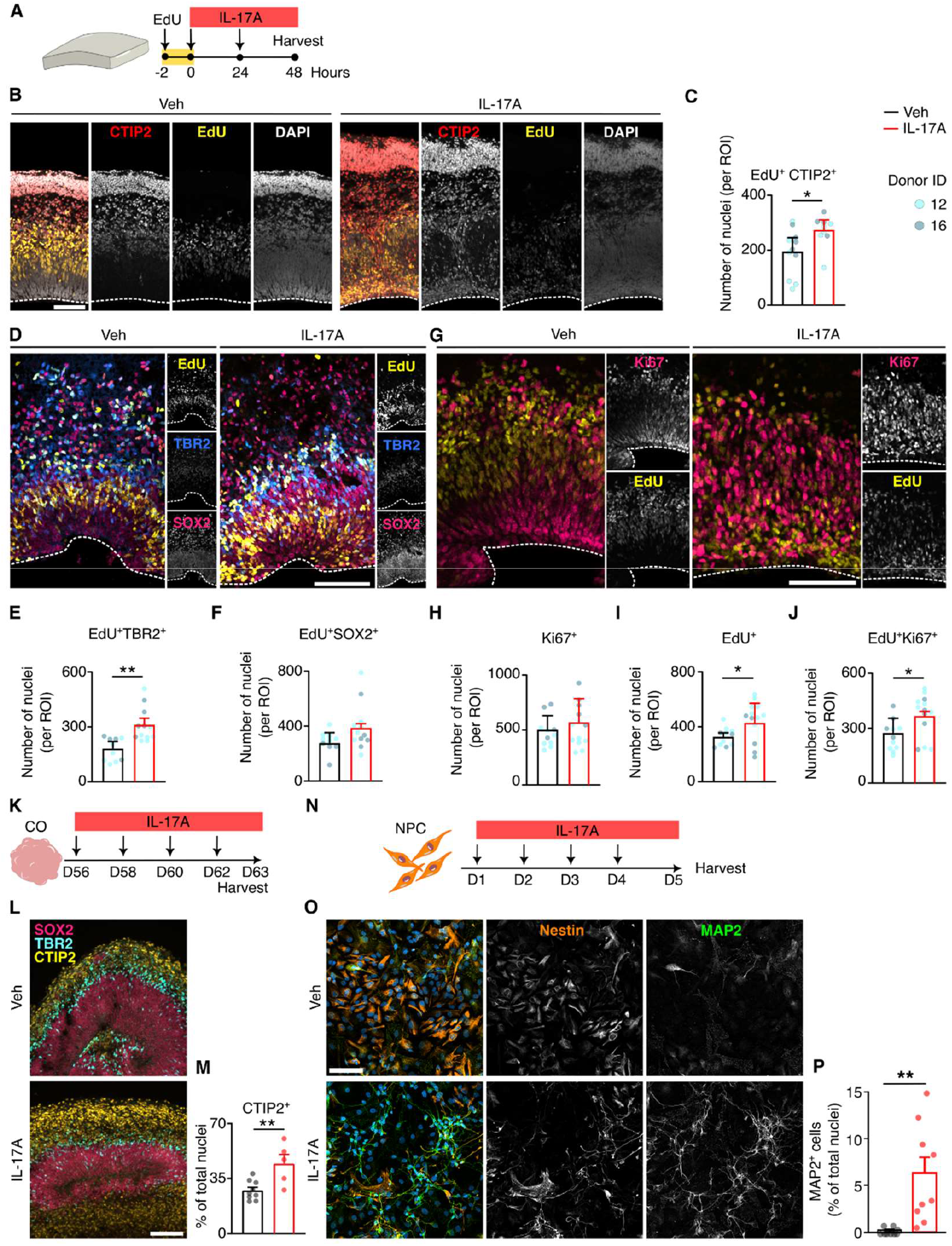
IL-17A affects the proliferation of NSCs in the developing human prefrontal cortex. **(A)** Schematic of the experimental paradigm for IL-17A and EdU administration to cerebroids. **(B** and **C)** Representative fluorescence images and quantification of EdU^+^ CTIP2^+^ neurons in vehicle- and IL-17A-treated cerebroids. (n = 6 cerebroids per group from 2 donors). **(D)** Representative fluorescence images showing EdU incorporation in neural stem cells (NSCs, SOX2^+^) and intermediate progenitor cells (IPCs, TBR2^+^) in vehicle- and IL-17A-treated cerebroids. **(E** and **F)** Quantification of EdU^+^ TBR2^+^ IPCs (E) and EdU^+^ SOX2^+^ NSCs (F) in vehicle- and IL-17A-treated cerebroids. (n = 6 cerebroids from 2 donors in each group). **(G)** Representative fluorescence images showing EdU incorporation and cell cycle marker Ki67 in vehicle- and IL-17A-treated cerebroids. **(H** to **J)** Quantification of the number of Ki67^+^ cells (H), EdU^+^ cells (I), and EdU/Ki67 double-positive cells (J) in vehicle- and IL-17A-treated cerebroids. (n = 6 cerebroids from 2 donors in each group). **(K)** Schematic of the experimental paradigm for IL-17A treatment of cortical organoids (COs) generated from human induced pluripotent stem cells (iPSCs). **(L)** Representative fluorescence image of COs stained for SOX2 (NSCs), TBR2 (IPCs), and CTIP2 (neurons) after IL-17A treatment. **(M)** Quantification of CTIP2^+^ cells in vehicle-(n = 9) and IL-17A-treated COs (n = 5). **(N)** Schematic of the experimental paradigm for IL-17A treatment of iPSC-derived neural precursor cells (NPCs). **(O)** Representative fluorescence images of Nestin+ (NPCs) and MAP2^+^ (neurons) in vehicle- and IL-17A-treated NPCs. **(P)** Quantification of MAP2^+^ cells in vehicle-and IL-17A-treated NPCs. (n = 3 independent experiments). Data is represented as mean ± SEM (*p < 0.05, **p < 0.01). Two-way ANOVA for treatment-group and donor in (C), (E), (F) and (H) to (J), and Student’s t-test for (M) and (P). Scale bar, 100 μm.

### Remodeling of ECM induced by IL-17A

To uncover the molecular mechanisms underlying abnormal cortical development, we performed bulk RNA-sequencing on whole cerebroids, as well as on three isolated cell populations: microglia/BAM, neuron-lineage committed cells (NLCCs), and NSCs **(Fig. 4A)** (*27*). The cellular identities of these isolated populations were confirmed by expression levels of marker genes **(fig. S8A)**. When comparing differentially expressed genes (DEGs) between vehicle-treated control and IL-17A-treated cerebroids, we identified five common categories of DEGs, including pathways related to epigenetic regulation/Sin-HDAC complex, TNF/NF-kB signaling, ECM and ECM-related glycosylation, energy metabolism, and WNT signaling **(Fig. 4, B to E, fig. S8B)**. While each population exhibited distinct Reactome pathway alterations, ECM-related pathways were consistently enriched across all cell types analyzed **(fig. S8B, fig. S9, A and B, fig. S10, fig. S11)**. Specifically, in NLCCs, IL-17A upregulated genes involved in ECM remodeling processes, including *THBS2, THBS1, COL4A2, COL11A2, LAMA2, DSEL*, and *COL9A2*, which are associated with integrin cell surface interactions, ECM degradation, collagen degradation, and collagen chain trimerization in Reactome enrichment analysis **(Fig. 4F)**. Mounting evidence shows that mitochondrial dynamics significantly influence cellular metabolism and shape the behavior of NSCs during brain development (*28*). Notably, mitochondria-related metabolic pathways were uniquely enriched in the Reactome pathway analysis of NSCs **(fig. S11)**, suggesting that altered mitochondria dynamics induced by IL-17A may contribute to the increased neurogenesis.

**Fig 4.**
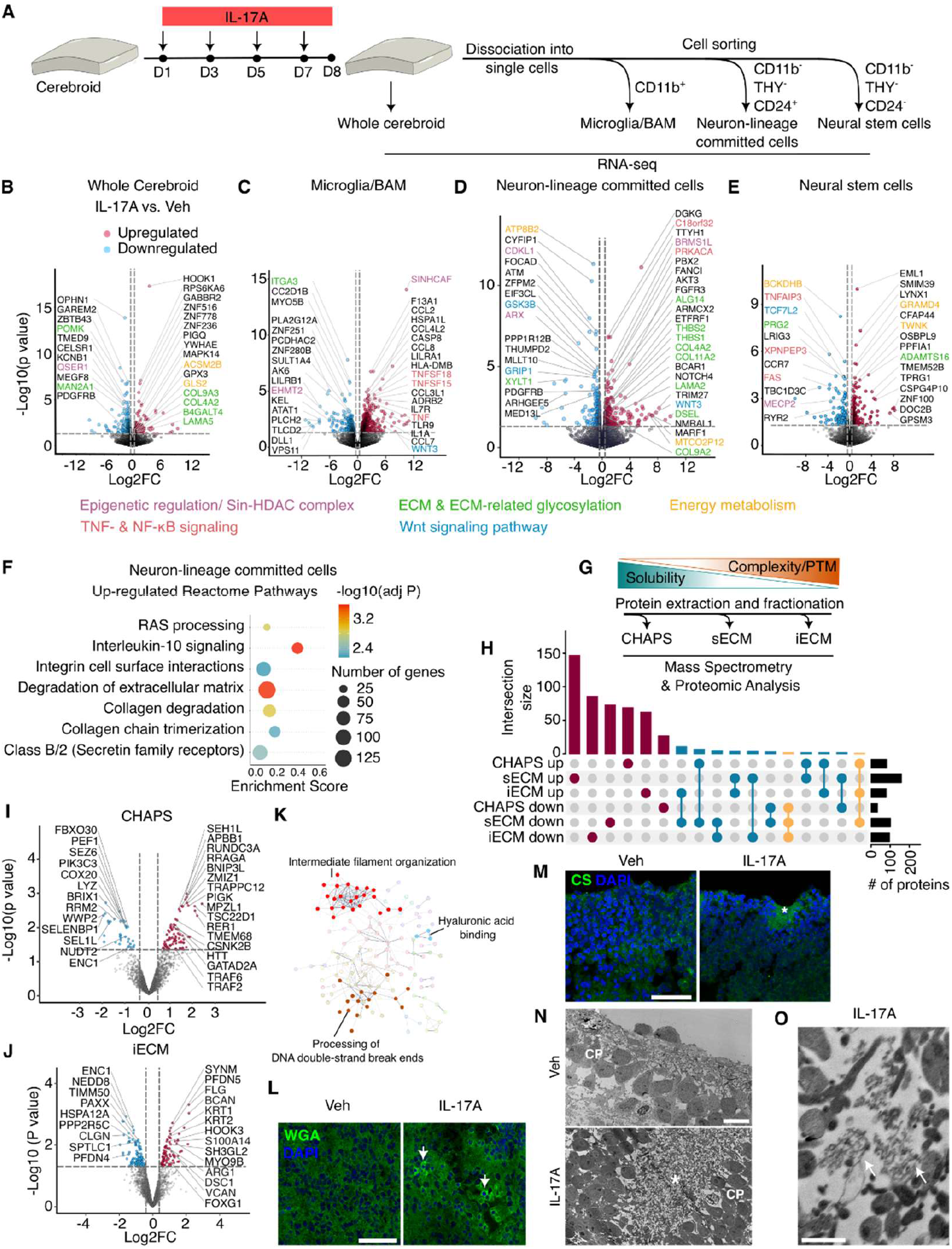
Transcriptional and proteomic changes in cerebroids following IL-17A treatment. **(A)** Schematic of the experimental design for RNA sequencing in bulk cerebroid tissue, microglia (CD11b^+^), neuron-lineage committed cells (NLCCs) (CD11b^-^, THY^-^ and CD24^+^), and neural stem cells (NSCs) (CD11b^-^, THY^-^ and CD24^-^). **(B** to **E)** Volcano plots of differentially expressed genes (DEGs) in IL-17A-versus vehicle-treated cerebroids for bulk tissue (B), microglia (C), NLCCs (D), and NSCs (E). Red denotes upregulated, blue denotes downregulated genes (n = 3 donors). **(F)** Bubble plot of enriched Reactome pathways from protein sub-network-based gene set enrichment analysis, showing upregulated genes in NLCCs after IL-17A treatment. Bubble color indicates FDR-adjusted p-value (Benjamini-Hochberg method), and size reflects the number of associated genes. **(G)** Schematic of protein extraction for mass spectrometry. Cellular proteins (CHAPS fraction), soluble extracellular matrix (sECM), and insoluble extracellular matrix (iECM) fractions were isolated and analyzed separately. **(H)** UpSet plot showing the number of differentially abundant proteins across three fractions (CHAPS, sECM, iECM) in IL-17A-versus vehicle-treated cerebroids (n = 18 samples: 2 experimental groups, 3 donors, 3 fractions per donor). **(I** and **J)** Volcano plots of differentially abundant proteins in the CHAPS (I) and iECM (J) fractions following IL-17A treatment. **(K)** Protein-protein interaction network of differentially abundant proteins in the iECM fraction, visualized using STRING clustering. Colors represent distinct sub-networks generated by k-means clustering. **(L)** Representative fluorescence images of wheat germ agglutinin (WGA) staining in vehicle- and IL-17A-treated cerebroids. Arrows indicate stronger WGA staining in IL-17A-treated cerebroids. Scale bar, 50 μm. **(M)** Representative fluorescence images of chondroitin sulfate (CS) staining, showing enhanced CS signal intensity at the cortical plate folding site in IL-17A-treated cerebroids. Scale bar, 50 μm. **(N** and **O)** TEM images of cortical plate regions showing areas of folding (asterisk) with abundant ECM in IL-17A-treated cerebroids (N). Scale bar, 5 μm. Higher magnification of glycosaminoglycan-containing proteoglycans (indicated by arrows) in the ECM (O). Scale bar, 500 nm.

Moreover, many DEGs in IL-17A-treated cerebroids were associated with neurodevelopmental and neurological disorders **(fig. S12)**, suggesting that IL-17A may dysregulate shared pathways involved in these conditions. We further conducted a meta-analysis of transcriptomic data from post-mortem brains of individuals with ASD **(fig. S13)** (*29-31*). Enrichment analysis revealed that many upregulated genes are linked to ECM components, reinforcing the significance of the ECM in the pathogenesis of NDDs.

To examine the ECM alterations induced by IL-17A, we performed mass spectrometry and proteomic analysis from three different protein fractions based on the solubility and complexity of proteins: CHAPS fraction, containing cellular proteins, the soluble ECM (sECM) fraction and insoluble ECM (iECM) fraction **(Fig. 4G and fig. S14A)**. We observed more than four hundred differentially abundant sECM and iECM proteins in IL-17A treated cerebroids. **(Fig. 4, H to J and fig. S14B)**. Protein-protein interaction network analysis of differentially abundant proteins in the CHAPS fraction of IL-17A-treated cerebroids revealed host-pathogen interaction of virus and IL-17A mediated signaling pathways, highlighting the activation of immune responses by IL-17A **(fig. S15)**. In the iECM fraction of IL-17A-treated cerebroids, two key chondroitin sulfate proteoglycans, brevican (BCAN) and versican (VCAN), were upregulated **(Fig. 4J)**. BCAN is a major proteoglycan in the brain’s ECM, playing critical roles in neural development, synaptic plasticity, and neuronal stability (*32-35*). VCAN contributes to the structural integrity of the developing ECM and is involved in cell proliferation, migration, and synapse formation (*36-39*). Both BCAN and VCAN bind to hyaluronic acid, a key ECM component that regulates cortical folding, cell migration, proliferation, axon guidance, and synapse formation (*12, 40*). Hyaluronic acid binding was also highlighted in the protein-protein interaction network in the iECM fraction of IL-17A-treated cerebroids, along with intermediate filament organization and processing of DNA double-strand break ends **(Fig. 4K)**.

The altered proteoglycan expression was visualized using wheat germ agglutinin (WGA), showing increased perineuronal net immunoreactivity in IL-17A-treated cerebroids **(Fig. 4L and fig. S16A)**. Additionally, chondroitin sulfate immunoreactivity was elevated at the sites of cortical folding in IL-17A-treated cerebroids **(Fig. 4M)**, aligning with evidence implicating the ECM in cortical folding (*41, 42*). TEM further confirmed the presence of increased glycosaminoglycan-containing proteoglycans at the folding sites at the ultrastructural level **(Fig. 4, N and O)**. These findings suggest a strong association between altered ECM composition and IL-17A-induced cortical folding. We also observed that expression of BCAN was heightened in ventricular zones of IL-17A treated cerebroids **(fig. S16B)**. These alterations in ECM components are particularly significant considering recent findings showing increased BCAN and VCAN expression in postmortem brains of children with ASD, with BCAN specifically elevated in astrocytes(*43*).

### Reversed abnormal cortical development by Parthenolide

To further investigate the molecular pathways mediating the abnormal cortical development induced by IL-17A during MIA, we identified key transcriptional regulators modulating the expression of DEGs from our RNA sequencing analysis. Among the many transcriptional regulators, five - HDAC1, NFKB1, REL, RELA, and SPI1 - were found to be shared across at least two different cell types **(Fig. 5A)**. Parthenolide (PTL) is a known inhibitor of the NF-κB pathway, through its suppression of IκB kinase activity, and an inhibitor of HDAC1 (*44, 45*). To confirm the involvement of these pathways in the abnormal cortical development phenotype, cerebroids were treated with PTL **(Fig. 5B)**. While PTL alone did not induce any observable phenotype, it reversed the cortical folding and thickness abnormalities across different cortical layers in IL-17A-treated cerebroids **(Fig. 5, C to I)**. Furthermore, PTL treatment mitigated the excessive neurogenesis, as evidenced by a reduction in the number of CTIP2^+^, SATB2^+^ and NeuN^+^ neurons in IL-17A-treated cerebroids **(Fig. 5, J and K)**. Given the upregulation of BCAN in the ventricular zone of IL-17A-treated cerebroids **(fig. S16B)**, we investigated whether PTL could reverse this effect as well. Following PTL treatment, BCAN expression in the ventricular zone was decreased, reaching levels similar to those observed in the vehicle group **(Fig. 5L)**. These data demonstrate that the NF-κB and HDAC1pathways play a crucial role in IL-17A-mediated abnormal cortical development during MIA.

**Fig 5.**
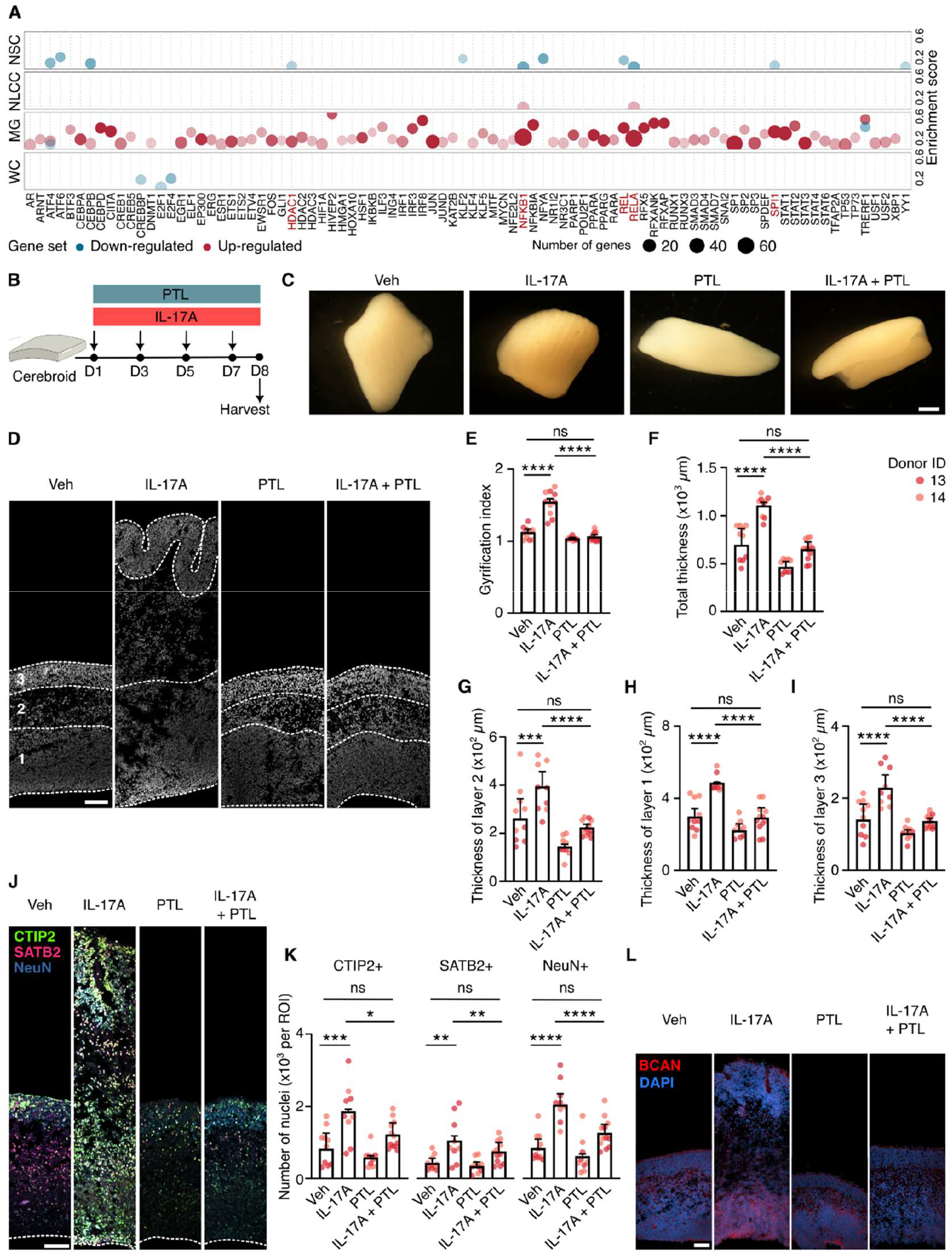
Parthenolide (PTL) reverses IL-17A-induced effects on cortical folding and neurogenesis in cerebroids. **(A)** Bubble plot of significant human transcription factors (TFs) from the TRRUST database regulating DEGs in IL-17A-treated whole cerebroids (WC), microglia (MG), NLCCs, and NSCs. Bubble color indicates direction of change, and size represents the number of regulated DEGs (FDR-adjusted p < 0.05). Analysis was performed with EnrichR. **(B)** Schematic illustration of PTL and IL-17A administration to cerebroid cultures. **(C)** Representative images of cerebroids treated with vehicle (Veh), IL-17A, PTL, and IL-17A + PTL. **(D)** DAPI-stained sections of cerebroids showing cortical architecture changes across treatment groups. Layer 1: ventricular and subventricular zones (VZ-SVZ), Layer 2: intermediate zone and subplate (IZ-SP), and Layer 3: cortical plate (CP). **(E)** Quantification of the gyrification index across treatment groups. **(F** to **I)** Quantification of cortical layer thickness in different treatment groups. **(J** and **K)** Representative fluorescence images (J) and quantification of CTIP2- and SATB2-expressing neurons (K) in cerebroids across different treatment groups. **(L)** Representative fluorescence images of ECM component brevican (BCAN) across treatment groups. Data are shown as mean ± SEM (*p < 0.05, **p < 0.01, ***p < 0.001, ****p < 0.0001, two-way ANOVA for treatment group and donor and Tukey’s multiple comparisons test for inter-group differences). N = 5–6 cerebroids per group, from 2 donors. Scale bars: 1000 μm (C), 100 μm [(D), (J), (L)].

## Discussion

In this study, we developed a robust high-throughput 3D ex-vivo culture system, cerebroids, which faithfully preserve key components of cortical development such as cellular architecture, apical-basal polarity, and essential immune, vascular, and ECM elements. The preservation of cellular integrity, heterogeneity, and ECM niche within cerebroids makes this model uniquely suited for investigating normal developmental processes and cell-cell interactions, as well as disruptions caused by environmental factors.

Utilizing the cerebroid, we demonstrated that IL-17A induces cortical abnormalities associated with MIA, including premature cortical folding, increased thickness, and accelerated neuronal maturation, driven by enhanced neurogenesis in NSCs. Modeling MIA in cerebroids has revealed clinically relevant cellular and molecular phenotypes associated with NDDs, such as microglia-driven immune responses, structural cortical changes, an increased number of excitatory neurons, and alterations in the ECM (*20-23*). Furthermore, this model offers a unique opportunity to dissect the molecular and mechanistic basis of these abnormalities during early development, a stage that is typically inaccessible. Our transcriptomic and proteomic analyses revealed that IL-17A remodels the ECM, with contributions from various cortical cell types. While the ECM is known to play important role in various developmental processes (*12, 35*), the specific contributions of different cell types to ECM composition and organization, and how they uniquely remodel it in response to environmental insults such as MIA, remain poorly understood. The cell-type-specific alterations in the ECM caused by IL-17A remains a critical avenue for future research. Additionally, PTL treatment successfully reversed IL-17A-induced cortical folding and neurogenesis abnormalities, implicating the involvement of NF-κB and/or HDAC1 pathways. Further research is required to clarify the specific roles of these pathways and their downstream mechanisms in ECM remodelling, as well as their contributions to cortical alterations.

Epidemiological, clinical, and animal studies have shown that males are more vulnerable to NDDs associated with MIA, both in terms of prevalence and symptom manifestation (*46*). Sex differences in response to MIA are likely influenced by a complex interplay of hormonal, placental, and microglia-mediated inflammation (*47*), though the precise mechanisms remain unclear. Due to limited tissue availability, sex differences were not explored in this study and further research is needed to address this gap. In this study we cultured cerebroids for 15 days at most. A recent study demonstrated that fetal brain organoids can be maintained in long-term culture through continuous tissue fragmentation (*48*). While cerebroids provide a robust platform for investigating the acute effects of environmental insults, future efforts should focus on overcoming the challenges of long-term culture, ensuring the preservation of normal developmental trajectory, cytoarchitecture, and cellular integrity. Achieving this will enable more comprehensive, long-term investigations into the impact of environmental factors on cortical development. In conclusion, we have established cerebroids as a robust, physiologically relevant model for interrogating complex neurodevelopmental processes. We have also demonstrated their application as a platform to study IL-17A-mediated disruptions in cortical development during MIA. In addition to incorporating neuronal and glial elements, cerebroids also integrate immune and vascular components, along with the ECM. This physiologically relevant complexity offers a unique platform to explore cell-cell and cell-ECM interactions across diverse cell populations, providing valuable insights into healthy brain development and NDDs.

## Acknowledgements

We thank the Kang and Berg labs for their support and insightful discussions. We gratefully acknowledge the Center for Genome-Enabled Biology and Medicine, Proteomics Core Facility, and the Microscopy and Histology Core Facility staff at the University of Aberdeen for their technical assistance. We also extend our thanks to the members of the Fowler Lab for their invaluable support in the acquisition of fetal samples. Special thanks to Tara Sutherland for constructive feedback and sharing antibodies. Most importantly, we would like to express our deepest gratitude to the donors and the NHS staff involved in tissue collection, whose contributions made this research possible.

## Funding

M.Z.K.A was supported by a BBSRC (Eastbio) 4-year PhD studentship. D.H.G was funded by The Humane Research Trust Les Rhoades PhD scholarship. D.A.B was funded by BBSRC (BB/W008068/1) and Royal Society (RG\R2\232559). E.K was funded by The Academy of Medical Sciences Springboard (SBF007\100169).

The SAFeR study was funded by the UK Medical Research Council (MR/L010011/1 and MR/P011535/1) and the EU’s Horizon 2020 research and innovation programme under the Marie Skłodowska-Curie project PROTECTED (grant agreement number 722634) and FREIA and INITIALISE projects (grant agreement numbers 825100 and 101094099 respectively) to P.A.F.

## Author contributions

M.Z.K.A., D.A.B., and E.K. conceived the project and designed the experiments. M.Z.K.A., D.A.B., and E.K. wrote the manuscript with input from all authors. M.Z.K.A. performed the majority of the experiments. M.Y. contributed to optimizing the cerebroid and iPSC-derived microglia cultures. D.H.G. conducted part of the bioinformatic analyses and assisted with tissue collection. P.M. and S.V. were responsible for tissue sectioning, staining, and imaging. L.P. carried out tissue dissection and qPCR analysis. P.A.F. provided the human brain tissue samples used in the study.

## Competing interests

The authors have none to declare.

## Notes

### Competing Interest Statement

The authors have declared no competing interest.

## References

1. H. K. Kwon, G. B. Choi, J. R. Huh, Maternal inflammation and its ramifications on fetal neurodevelopment. Trends Immunol 43, 230–244 (2022).

2. V. X. Han, S. Patel, H. F. Jones, R. C. Dale, Maternal immune activation and neuroinflammation in human neurodevelopmental disorders. Nat Rev Neurol 17, 564–579 (2021).

3. H. O. Atladottir et al., Maternal infection requiring hospitalization during pregnancy and autism spectrum disorders. J Autism Dev Disord 40, 1423–1430 (2010).

4. A. G. Edlow et al., Sex-Specific Neurodevelopmental Outcomes Among Offspring of Mothers With SARS-CoV-2 Infection During Pregnancy. JAMA Netw Open 6, e234415 (2023).

5. B. J. S. Al-Haddad et al., Long-term Risk of Neuropsychiatric Disease After Exposure to Infection In Utero. JAMA Psychiatry 76, 594–602 (2019).

6. N. V. Malkova, C. Z. Yu, E. Y. Hsiao, M. J. Moore, P. H. Patterson, Maternal immune activation yields offspring displaying mouse versions of the three core symptoms of autism. Brain Behav Immun 26, 607–616 (2012).

7. G. B. Choi et al., The maternal interleukin-17a pathway in mice promotes autism-like phenotypes in offspring. Science 351, 933–939 (2016).

8. L. Ben-Reuven, O. Reiner, Dynamics of cortical progenitors and production of subcerebral neurons are altered in embryos of a maternal inflammation model for autism. Mol Psychiatry 26, 1535–1550 (2021).

9. K. W. Kelley, S. P. Pasca, Human brain organogenesis: Toward a cellular understanding of development and disease. Cell 185, 42–61 (2022).

10. Y. K. Adlakha, Human 3D brain organoids: steering the demolecularization of brain and neurological diseases. Cell Death Discov 9, 221 (2023).

11. M. G. Andrews, L. Subramanian, J. Salma, A. R. Kriegstein, How mechanisms of stem cell polarity shape the human cerebral cortex. Nat Rev Neurosci 23, 711–724 (2022).

12. K. R. Long, W. B. Huttner, The Role of the Extracellular Matrix in Neural Progenitor Cell Proliferation and Cortical Folding During Human Neocortex Development. Front Cell Neurosci 15, 804649 (2021).

13. M. E. Akintunde et al., Increased production of IL-17 in children with autism spectrum disorders and co-morbid asthma. J Neuroimmunol 286, 33–41 (2015).

14. K. L. Jones et al., Autism with intellectual disability is associated with increased levels of maternal cytokines and chemokines during gestation. Mol Psychiatry 22, 273–279 (2017).

15. M. Moaaz, S. Youssry, A. Elfatatry, M. A. El Rahman, Th17/Treg cells imbalance and their related cytokines (IL-17, IL-10 and TGF-beta) in children with autism spectrum disorder. J Neuroimmunol 337, 577071 (2019).

16. B. van der Zwaag et al., Gene-network analysis identifies susceptibility genes related to glycobiology in autism. PLoS One 4, e5324 (2009).

17. H. Wong, C. Hoeffer, Maternal IL-17A in autism. Exp Neurol 299, 228–240 (2018).

18. K. Yasumatsu et al., Bacterial-induced maternal interleukin-17A pathway promotes autistic-like behaviors in mouse offspring. Exp Anim 69, 250–260 (2020).

19. K. Sarieva et al., Human brain organoid model of maternal immune activation identifies radial glia cells as selectively vulnerable. Mol Psychiatry 28, 5077–5089 (2023).

20. E. Fombonne, B. Roge, J. Claverie, S. Courty, J. Fremolle, Microcephaly and macrocephaly in autism. J Autism Dev Disord 29, 113–119 (1999).

21. M. G. Butler et al., Subset of individuals with autism spectrum disorders and extreme macrocephaly associated with germline PTEN tumour suppressor gene mutations. J Med Genet 42, 318–321 (2005).

22. E. Courchesne et al., Neuron number and size in prefrontal cortex of children with autism. JAMA 306, 2001–2010 (2011).

23. C. W. Nordahl et al., Cortical folding abnormalities in autism revealed by surface-based morphometry. J Neurosci 27, 11725–11735 (2007).

24. Z. Molnar et al., New insights into the development of the human cerebral cortex. J Anat 235, 432–451 (2019).

25. E. C. Gilmore, C. A. Walsh, Genetic causes of microcephaly and lessons for neuronal development. Wiley Interdiscip Rev Dev Biol 2, 461–478 (2013).

26. B. E. L. Ostrem et al., Fetal brain response to maternal inflammation requires microglia. Development 151, (2024).

27. D. D. Liu et al., Purification and characterization of human neural stem and progenitor cells. Cell 186, 1179–1194 e1115 (2023).

28. V. Scandella, F. Petrelli, D. L. Moore, S. M. G. Braun, M. Knobloch, Neural stem cell metabolism revisited: a critical role for mitochondria. Trends Endocrinol Metab 34, 446–461 (2023).

29. N. N. Parikshak et al., Genome-wide changes in lncRNA, splicing, and regional gene expression patterns in autism. Nature 540, 423–427 (2016).

30. M. J. Gandal et al., Shared molecular neuropathology across major psychiatric disorders parallels polygenic overlap. Science 359, 693–697 (2018).

31. M. J. Gandal et al., Broad transcriptomic dysregulation occurs across the cerebral cortex in ASD. Nature 611, 532–539 (2022).

32. M. Blosa et al., The extracellular matrix molecule brevican is an integral component of the machinery mediating fast synaptic transmission at the calyx of Held. J Physiol 593, 4341–4360 (2015).

33. R. Frischknecht, C. I. Seidenbecher, Brevican: a key proteoglycan in the perisynaptic extracellular matrix of the brain. Int J Biochem Cell Biol 44, 1051–1054 (2012).

34. C. Gottschling, D. Wegrzyn, B. Denecke, A. Faissner, Elimination of the four extracellular matrix molecules tenascin-C, tenascin-R, brevican and neurocan alters the ratio of excitatory and inhibitory synapses. Sci Rep 9, 13939 (2019).

35. K. R. Long, W. B. Huttner, How the extracellular matrix shapes neural development. Open Biol 9, 180216 (2019).

36. F. Sotoodehnejadnematalahi, B. Burke, Structure, function and regulation of versican: the most abundant type of proteoglycan in the extracellular matrix. Acta Med Iran 51, 740–750 (2013).

37. M. Schmalfeldt, C. E. Bandtlow, M. T. Dours-Zimmermann, K. H. Winterhalter, D. R. Zimmermann, Brain derived versican V2 is a potent inhibitor of axonal growth. J Cell Sci 113 (Pt 5), 807–816 (2000).

38. H. Watanabe, Aggrecan and versican: two brothers close or apart. Am J Physiol Cell Physiol 322, C967–C976 (2022).

39. M. O. Politko et al., Multiple Irradiation Affects Cellular and Extracellular Components of the Mouse Brain Tissue and Adhesion and Proliferation of Glioblastoma Cells in Experimental System In Vivo. Int J Mol Sci 22, (2021).

40. R. Frischknecht, C. I. Seidenbecher, The crosstalk of hyaluronan-based extracellular matrix and synapses. Neuron Glia Biol 4, 249–257 (2008).

41. K. R. Long et al., Extracellular Matrix Components HAPLN1, Lumican, and Collagen I Cause Hyaluronic Acid-Dependent Folding of the Developing Human Neocortex. Neuron 99, 702–719 e706 (2018).

42. S. Amin, V. Borrell, The Extracellular Matrix in the Evolution of Cortical Development and Folding. Front Cell Dev Biol 8, 604448 (2020).

43. L. E. Rexrode et al., Molecular profiling of the hippocampus of children with autism spectrum disorder. Mol Psychiatry 29, 1968–1979 (2024).

44. B. H. Kwok, B. Koh, M. I. Ndubuisi, M. Elofsson, C. M. Crews, The anti-inflammatory natural product parthenolide from the medicinal herb Feverfew directly binds to and inhibits IkappaB kinase. Chem Biol 8, 759–766 (2001).

45. Y. N. Gopal, T. S. Arora, M. W. Van Dyke, Parthenolide specifically depletes histone deacetylase 1 protein and induces cell death through ataxia telangiectasia mutated. Chem Biol 14, 813–823 (2007).

46. S. Sal-Sarria, N. M. Conejo, H. Gonzalez-Pardo, Maternal immune activation and its multifaceted effects on learning and memory in rodent offspring: A systematic review. Neurosci Biobehav Rev 164, 105844 (2024).

47. H. C. Osman et al., Impact of maternal immune activation and sex on placental and fetal brain cytokine and gene expression profiles in a preclinical model of neurodevelopmental disorders. J Neuroinflammation 21, 118 (2024).

48. D. Hendriks et al., Human fetal brain self-organizes into long-term expanding organoids. Cell 187, 712–732 e738 (2024).

