## supplementary information for "IL-17A Alters Human Cortical Development in a 3D Ex Vivo Model of Maternal Immune Activation"

**The PDF file includes:**

Materials and Methods
Figs. S1 to S16
Tables S1-4
References 1-38

### Materials and Methods

#### Human fetal tissue collection

Fetal tissues from elective, normally progressing pregnancies were collected under the Scottish Advanced Fetal Research (SAFeR) study (NCT04613583) (*1*). The collection process is approved by one of the 12 Scottish National Health Service Research Ethics Committees (REC 15/NS/0123) and follows the Declaration of Helsinki guidelines. Women seeking elective medical termination of pregnancy were recruited with written informed consent by NHS Grampian research nurses, who operate independently from the research team. There was no alteration in patient treatment or care, and participants could withdraw from the study at any time up to the point the fetus was removed from the ward. The study includes only normally progressing pregnancies, as determined by ultrasound, from women over 16 years of age who speak English and covers gestational ages from 7 to 20 weeks. Grossly abnormal fetuses were excluded, and women experiencing significant distress were not approached. Termination was carried out using RU-486 (Mifepristone) treatment (200 mg) followed by prostaglandin-induced delivery, as reported previously (*2*). Gestational age was confirmed by ultrasound and foot length measurement, and various maternal and fetal data were recorded. For sample collection, fetuses are transported to the laboratory within 30 minutes of delivery, typically intact, and are weighed, measured for crown-rump length (CRL), and sexed by morphology and PCR detection of sex chromosome-specific genes (ZFX and SRY) or the absence of the Y chromosome. Brain tissues from 16 donors of 11 to 14 weeks of gestation collected between 11/05/2022 and 22/04/2024 were included in this study (see **table S1**).

#### Brain dissection and tissue culture

Whole brain tissue was transferred to cold Hanks' balanced salt solution (HBSS) (Gibco, 14175095) on ice for dissection. The pia matter was carefully removed. Full thickness approximately $10mm^2$ size pieces of right dorso-lateral prefrontal cortical (DLPFC) tissue were dissected and plated in 6-well tissue culture plates (16-24 cerebroids per well). Dissection time was kept to a minimum (~30 minutes) and cerebroids were immediately transferred to the fresh culture medium (4 mL per well) containing a 1:1 mixture (v/v) of Dulbecco's Modified Eagle Medium/Nutrient Mixture F-12 (DMEM/F-12, Gibco, 11330-032) and Neurobasal medium (Gibco, 21103-049), supplemented with 1X Glutamax (Gibco, 35050-038), 1X MEM Non-essential amino acids solution (Gibco, 11140-050), 1X serum free B27 (Gibco, 17504-044), 1X N-2 (Gibco, 17502-048), 1X 2-Mercaptoethanol (Gibco, 21985-023), 2.5 μg/ml human recombinant Insulin (Gibco, 12585-014), and 1X Antibiotic-Antimycotic (Gibco, 15240-096). Cerebroids were cultured in a free-floating manner on a $CO_2$-resistent orbital shaker (Fisher Scientific, 15351105) at a rotational rate of 120/min in a humidified incubator (37°C and 5% $CO_2$). Culture medium was changed on alternate days.

#### Cell culture

The human induced pluripotent stem cell (iPSCs) line KOLF2.1J (Wellcome Sanger Institute, RRID:CVCL_B5P3) was used to generate cortical organoids and iPSC-derived microglia like cells (iMGLs). Human iPSCs were culture in mTeSR plus medium containing mTeSR plus basal medium (StemCell Technologies, 100-0274), mTeSR plus 5X supplement (StemCell Technologies, 100-0275), and 1X Penicillin-Streptomycin (Gibco, 15140-122).

Human iPSC line SCTi003-A—derived neural progenitor cells (NPCs) (Stemcell Technologies, 200-0620) were cultured in STEMdiff neural progenitor medium (Stemcell technologies, 05833) supplemented with 1X Penicillin-Streptomycin according to the supplier's instructions. Media was fully changed daily and cells were passaged every 7-10 days using ACCUTASE (StemCell technologies, 7920) and media supplementation with 10μM Y-27632 (Dihydrochloride) ROCK inhibitor (StemCell Technologies, 72304) for first 16-24 hours of plating the cells.
NPCs were differentiated into forebrain neurons using STEMdiff Forebrain Neuron Differentiation Medium (StemCell technologies, 08600) in Laminin/Poly-L-Ornithine (Sigma-Aldrich, LPLO001) coated 12-well plates. Media was changed daily. After 5-6 days of neuronal differentiation, cells were detached using ACCUTASE and replated on Laminin/Poly-L-Ornithine coated 12-well plates at a seeding density of $4 \times 10^4$ - $6 \times 10^4$ cells/cm$^2$. Cells were cultured in STEMdiff Forebrain Neuron Maturation Medium (StemCell technologies, 08605) and full media change was done every 2 days.
Human iPSC-derived microglia like cells (iMGLs) were generated from Kolf2.1J cells using a previously published protocol (*3*).

**iPSC-derived cortical organoid culture**
Human KOLF2.1J iPSC-derived cortical organoids were generated using previously published protocol (*4, 5*) with few modifications. Briefly, embryoid bodies (EB) were generated from $5 \times 10^4$ iPSCs/EB. Cells were suspended in mTeSR plus supplemented with 1:2000 ROCK inhibitor and added to wells of an ultra-low attachment 96 U-bottom well plate (Corning, 7007). Plates were centrifuged at 300*xg* for 5min at 4°C. Cells were then incubated at 37°C and 5% $CO_2$ for 48 hours. EBs were collected and cultured in DMEM/F-12 supplemented with 20% knock out serum replacement (KOSR) (Gibco, 10828-028), 1X Glutamax, 1X MEM Non-essential amino acids solution, 1X 2-Mercaptoethanol, 1X Penicillin/Streptomycin (Gibco, 15140-122), 1 μM LDN (StemCell Technologies, 72147), and 5 μM SB, 1:1000 0.2% Heparin Solution (StemCell Technologies, 7980) for 7 days on an orbital shaker at 120 rpm, 37°C and 5% $CO_2$. Media was changed every other day and was supplemented with 1:2000 ROCK inhibitor for first 48 hours. Forebrain patterning of EBs were done in DMEM/F12 supplemented with 1X Glutamax, 1X MEM Non-essential amino acids solution, 1X N-2, 1X 2-Mercaptoethanol (Gibco, 21985-023), 1X Penicillin/Streptomycin, and 3μM CHIR (StemCell Technologies, 72054). After 7 days, EBs were cultured on orbital shaker at 120 rpm, 37°C and 5% $CO_2$ in a medium containing a 1:1 mixture (v/v) of DMEM/F-12 and Neurobasal medium, supplemented with 1X Glutamax, 1X MEM Non-essential amino acids solution, 1X serum free B27, 1X N-2, 1X 2-Mercaptoethanol, 2.5 μg/ml human recombinant Insulin, and 1X Penicillin/Streptomycin. Two-third media change was done every other day.

**IL-17A treatment**
Maternal immune activation was modelled by treating cerebroids with 100ng/mL human recombinant IL-17A (Peprotech, 200-17) every other day for a week **(Fig. 2A)**. The dose was chosen based on the prior in vivo (*6-10*) and 2D (*7, 11, 12*) and 3D (*8, 13, 14*) in vitro studies. The dose of IL-17A used in prior studies has been quite variable ranging from 10 ng/mL to 500 ng/mL in 2D and 25 ng/mL to 500 ng/mL of culture medium in 3D in vitro cultures. We tested

both 100 ng/mL and 200 ng/mL dose of IL-17A for cerebroids and found comparable changes in cortical thickness and folding phenotype (data not shown) and opted to add 100 ng/ml of IL-17A to the cerebroid culture every 48 hours. For both vehicle- and IL17A-treated groups, two third of the medium was replaced with fresh medium containing vehicle or IL17A.
2D cultured NPCs were treated with 12.5 ng/mL of IL-17A daily for four days **(Fig. 3P)**. NPC-derived forebrain neurons were treated with 12.5 ng/mL of IL-17A every 2 days for 1 week **(fig. S6F)**.

**Parthenolide treatment**
(-)-Parthenolide (PTL) (Selleckchem, S2341) was dissolved in DMSO to constitute a 12.5mM stock solution. Cerebroids were treated with 25 µM PTL in cerebroid culture medium on alternative days for a week with and without IL-17A treatment **(Fig. 5A)**. 0.2% DMSO was used as vehicle control.

**Tissue preparation and immunofluorescent staining**
Cerebroids were fixed in 4% Paraformaldehyde (PFA) in Phosphate Buffered Saline (PBS) for 60 min at room temperature and cryoprotected with 30% sucrose overnight at 4°C. Fetal brain tissue was fixed in 4% PFA in PBS overnight and cryoprotected sequentially with 15% and 30% sucrose at 4°C. Cortical organoids were fixed in 4% PFA in PBS for 30 min at room temperature and cryoprotected as described for cerebroids. Cerebroids and fetal brain tissue were embedded in OCT Embedding matrix (VWR Chemicals, 361603E), frozen on dry ice, sectioned at -20°C in a cryostat (Leica Biosystems, CM1850) perpendicular to the ventricular surface at 20µm thickness, and stored at -20°C. Frozen slides were gradually equilibrated to room temperature before immunostaining. For each set of staining, a minimum of 3-4 sections 160-200µm apart were included per cerebroid. Tissue sections were washed and permeabilized with TBS+ (0.05M Tris Buffered Saline, 0.1% Triton-X, pH 9.0) for 5 min three times followed by the antigen retrieval, when necessary, with 1X Target Retrieval Solution (Dako Agilent, S169984-2) at 88-90°C for 15 min in a plastic coplin staining jar (Epredia, 194). Tissue sections were washed again three times with TBS+ for 5 min and were blocked with the blocking solution TBS plus (containing 10% donkey serum in TBS with 0.1% Triton-X) for 1 hour at room temperature. Tissue sections were incubated with primary antibodies diluted in TBS with 0.1% Triton-X and 3% donkey serum overnight at 4°C in an immunostaining moisture chamber (Agar Scientific, AGL4555). Longer incubation time (48-72 hours) was need for primary staining with anti-HOPX antibodies. After washing three times with TBS+, tissues were incubated with species-specific secondary antibodies and DAPI (1:5000) diluted in TBS+ with 3% donkey serum for 2 hours at room temperature. Slides were washed three times with TBS+ and mounted. The details of primary and secondary antibodies used in this study are provided in **table S2 and 3**.
NPCs cultured on chamber slides were fixed with 4% PFA in PBS at room temperature for 15 minutes followed by washing with PBS three times. Immunostaining was performed as described above with only differences being no antigen retrieval and shorter incubation time with secondary antibodies (1 hour).
For WGA staining of fixed tissue, cerebroid sections were washed with PBS without permeabilization agents, incubated with Alexa Fluor 555 conjugated WGA (Invitrogen, 11590816) in PBS at 10 µg/mL for 30-60 minutes at room temperature. Slides were washed three times with PBS and nuclei were stained with DAPI.

**EdU pulse-chase experiments**
For 5'-Ethynyl-2'-deoxyuridine (EdU) pulse-chase experiments, 10 µM EdU (Invitrogen, A10044) was added to culture medium and incubated at 37°C on a shaker for 2 hours. After 2 hours, the media was removed, cerebroids were washed three times with DPBS and fresh medium without EdU was added **(Fig. 1I and Fig. 3A)**.
EdU detection was done after incubation with the secondary antibodies using Click-iT EdU cell proliferation kit for imaging, Alexa Fluor 647 dye (Invitrogen, C10340) according to the manufacturer's instructions.

**Genetic labelling of radial glia cells using adenovirus**
The recombinant human Adenovirus Type5 (dE1/E3) expressing enhanced green fluorescent protein (eGFP) (Vector Biolabs, 1060) or red fluorescent protein (RFP) (Vector Biolabs, 1660) under CMV promoter was used to selectively label apical radial glia cells in the cerebroids. ~15-20 µL of DMEM/F12 containing $1x10^7$ Plaque Forming Units (PFU)/µL Adenovirus was added directly on the ventricular surface of the cerebroids after removing the culture medium. Cerebroids were incubated for 20-30 minutes at 37°C and 5% $CO_2$ without agitation. Cerebroids were then washed three times with DPBS and cultured in the fresh medium.

**Calcium imaging**
Neurons were incubated with 1:1 v/v 10 µM Fluor-4-AM (Fisher Scientific, 11504786) and Pluronic acid (Fisher Scientific, P3000MP) in the culture medium for 30 minutes in the incubator. Cells were washed three times with HBSS without calcium and magnesium three times and then incubated with HBSS containing calcium and magnesium for 20 minutes at room temperature. Cells were imaged on EVOS M5000 imaging system (Thermo Fisher, AMF5000SV) with EVOS onstage incubator (Thermo Fisher, AMC2000) with the following settings: 37°C, 5% $CO_2$, and 80% humidity.
Images were analyzed using ImageJ. Movement artifacts were removed using Linear Stack Alignment with SIFT plugin and photo bleach correction performed using the exponential bleach correction function. Circular ROIs were drawn for individual neurons and fluorescence intensities were measured across the stack. ΔF/Fmin were calculated and plotted in GraphPad Prism. Heatmaps were drawn in R using ComplexHeatmap package v2.20.0 (https://bioconductor.org/packages/release/bioc/html/ComplexHeatmap.html).

**Immunofluorescence image acquisition and image analysis**
Images were acquired on a laser scanning confocal microscope (Carl Zeiss LSM880) using 20X, 40X (oil immersion) and 63X (oil immersion) objectives. Uniform laser settings were used across experimental groups where intensity-based analysis was planned. For larger tissue sections, tiled images were acquired to include all the layers (VZ to CP) and stitched automatically. For uniformity, images were obtained from the central $2/3^{rd}$ of the cerebroid sections. At least two to three sections were imaged for each cerebroid.
Images were analyzed using Fiji ImageJ software. To achieve consistent and reproducible results, we applied a uniform image acquisition, pre-processing, and analysis pipeline to count cells across the experimental groups for a given set of staining. Rectangular regions of interests (ROIs) were drawn with the base of the rectangle parallel to the ventricular surface and length

covering the whole thickness of the cerebroid section to include all layers **(fig. S7, A and B)**. The width of these ROIs was kept constant for comparison across experimental groups and replicates. For quantification of nuclei positive for nuclear markers (e.g. SOX2 and CTIP2), images were segmented through ImageJ interactive watershed plugin version 1.2.1 (https://imagej.net/plugins/interactive-watershed) and binary masks were generated. Uniform settings (seed dynamics, threshold intensity, and peak flooding) were used for segmenting nuclei across experimental groups for each set of staining. Number of binarized nuclei were quantified using the analyze particle function in ImageJ. For all quantitative analysis, images were blinded before quantification.

No pre-processing was performed for intensity-based quantification analysis **(Fig. 2, M and N)**. For nuclear fluorescence intensity measurement of different markers, we first segmented all nuclei from DAPI images using interactive watershed plugin version 1.2.1 with the following settings (seed dynamics 8-10, intensity threshold 20, and 100% peak flooding) **(fig. S7C)**. Following the segmentation, individual masks were generated for these nuclei. We measured the size and shape descriptors for these nuclei along with the X and Y-axis spatial coordinates of their centers. We then applied these masks to the 8-bit images of various markers (e.g. CTIP2, SATB2, and NeuN) and measured the fluorescence intensities (mean, mode, min, max, integrated density, and raw integrated intensity) for these individual nuclei using multi-measure function in ImageJ. Nuclei less than 8 or greater than 200 µm in size were filtered out. Fluorescence intensities were corrected for background noise by subtracting average background intensities.

Distance along the Y-axis was normalized for each individual image (0 being the ventricular surface and 1 being the pial surface) before combining data across multiple images or making multiple comparisons given the heterogeneous thickness of the cerebroids.

**Measurement of cortical folding**

Cortical folding was quantified by measuring gyrification index (GI) **(Fig. 2F)** on low magnification (4X or 10X) DAPI images of cerebroids.

Gyrification index was calculated as follows:

Gyrification index = Length of fiducial (inner) surface/ Length of (outer) convex hull

**Measurement of layer thickness**

Total thickness of a cerebroid was measured as an average of three vertical lines drawn perpendicular to the ventricular surface through the central part of the DAPI stained images of a cerebroid. Three layers of cerebroids (Layers 1-3) were defined based on the cell densities (layer 1 and 3 being cell dense, layer 2 being cell-sparse) and staining of specific markers (SOX2 and TBR2 enriched layer 1, CTIP2 enriched layer 3) **(Fig. 2E, fig. S6D, and fig. S7B)**. Thickness of each of these layers in the center of cerebroids was measured as average length of three vertical lines drawn along the layers perpendicular to the ventricular surface.

**Electron microscopy**

Tissue was fixed in 2% Glutaraldehyde (TAAB UK, G010) & 4% Formaldehyde (Fisher Scientific, 28909) in 0.1M Cacodylate (TAAB, S007) buffer for 48 hours at 4°C and stored in 0.1M Cacodylate buffer at 4°C till further processing. Samples were postfixed with 1% Osmium tetroxide (TAAB O021) and 1.5% Potassium Ferrocyanide (Sigma P-3289) in 0.1M Cacodylate buffer, washed and sequentially dehydrated with 35%, 50%, 70%, and 95% ethanol followed by acetone. This was followed by Spurr's infiltration and embedding in Spurr's resin for 48 hours at 60°C. 90nm thick sections were prepared using a diamond knife (Diatome Ltd, Switzerland) onto formvar/carbon coated Copper grids (EMResolutions, UK) using a Leica UC6 ultramicrotome and contrast stained with Uranyless (Labtech, UK) and Lead Citrate (Leica, UK) using a Leica AC20. Samples were viewed on the Transmission Electron Microscopy JEM 1400 plus (JEOL UK) and images were captured using an AMT UltraVUE camera (AMT, USA). For scanning electron microscopy (SEM), cerebroids were cut with a fine blade into half and each half was attached to the stub with either ventricular zone or cortical surface facing up. SEM images were acquired on ZEISS EVO MA10 SEM.

**Wheat Germ Agglutinin (WGA) Staining**
For live staining with Alexa fluor 555 conjugated WGA (Invitrogen, 11590816), cerebroids were transferred to wells of 12-well plate containing WGA diluted in HBSS at 10µg/mL for 30 minutes at room temperature. Cerebroids were washed three times with HBSS, imaged, and transferred back to the fresh culture medium.

**Assessment of the basal inflammation level of the donor fetal tissue**
Matching placental tissue was assessed for the expression of inflammatory markers as a proxy for the baseline in-utero inflammatory milieu of each fetus. Umbilical cords, fetal membranes, maternal decidua and any blood clots were removed from placentas prior to weighing. Each placental sample consisted of 6 full-thickness biopsies which were combined then snap-frozen and pulverized in liquid nitrogen to form a representative powder. Approximately 30 mg of the powdered tissue was used to extract DNA, RNA and protein according to manufacturer's guidelines (AllPrep DNA/RNA/Protein kit, 80004, QIAGEN UK). RNA was reverse transcribed into cDNA using the GoScript Reverse Transcriptase Kit (Promega, 2790). Representative pools of RNA (10-20 samples) were combined to make representative cDNA for standard curves. For quantitative real-time PCR (RT-qPCR), primers were designed to amplify the transcripts of inflammation markers C3, TNF, IL-6, IL-1B, NOS2 using Integrated DNA Technologies PCR designing web tool (https://www.idtdna.com/page) and are shown in table S4. Transcript expression was quantified by RT-qPCR (LightCycler 480 SYBR Green I Master, Roche) (Roche LightCycler 480). Samples were run in triplicate against a 6-point standard curve for each transcript (Absolute Quantification analysis) using parameters; 95 °C for 5 min, 45 cycles of 95 °C for 15 s, 63 °C for 15 s and 72 °C for 10 s followed by a melting curve. A selection of samples was repeated across plates to test for inter-plate variation. 5 candidate house-keeping genes (HKG) for placenta identified by previous studies (*15*) were tested for stabilities against fetal age, sex and maternal BMI. The geometric mean of the 3 most stable genes were used for normalization of the placenta transcript data.

**Tissue dissociation and single cell isolation**
Cerebroids were dissociated into single cells through enzymatic digestion and mechanical dissociation using a previously published protocol (*16*) with few modifications. Cerebroids was dissected into small pieces with a fine dissection scissor, followed by enzymatic digestion and mechanical dissociation with flame-polished Pasteur pipets in an enzyme mix containing Collagenase A (Millipore Sigma, 11088793001) and DNase I (Millipore Sigma, 10104159001) in 5% Fetal Bovine Serum (Fisher Scientific, 15383681), HBSS (without calcium and magnesium) and 10µM HEPES (Fisher Scientific, 15630-080). Myelin removal step with Percoll

method wasn't required. Duration of enzymatic digestion and mechanical dissociation was kept at 15 minutes. Uniform single cell suspension was obtained by filtering through a 70µm nylon filter (Corning, 431751).

**Immuno-magnetic cell sorting**

Microglia were isolated through immunomagnetic positive selection of $CD11b^{+}$ cell by using EasySep Human CD11b Positive Selection and Depletion Kit (Stemcell Technologies, 100-0742) and The Big Easy EasySep Magnet (Stemcell Technologies, 18001) according to the manufacturer's instructions. $CD24^{+}THY1^{-}CD11b^{-}$ and $CD24^{-}THY1^{-}CD11b^{-}$ cells representing NLCCs and NSCs, respectively, were isolated from CD11b depleted single cell population. $CD11b^{-}$ cells were subjected sequentially to immunomagnetic sorting with biotinylated anti-THY-1 (CD90) antibodies (Stemcell Technologies, 60045BT) and anti-CD24 antibodies (Miltenyi biotec, 130-098-902) using the EasySep Release Human Biotin Positive Selection Kit (Stemcell Technologies, 17653) and EasySep Magnet according to manufacturer's guidelines **(Fig. 4A)**.

**RNA extraction and RNA sequencing**

Total RNA was extracted from whole cerebroids and isolated cells using RNeasy mini kit (Qiagen, 74104) according to the manufacturer's instruction in an RNase free environment. For isolated microglia, cell lysis with RLT buffer was followed by an additional step of placing Eppendorf tubes in a magnetic rack (DynaMag-2, Invitrogen, 12321D) for 5 minutes and careful aspiration of clear cell lysate without antiCD11b magnetic beads. Residual genomic DNA was removed by on-column digestion with RNase-free DNase I (Qiagen, 79254). RNA quantity was measured on a Qubit Fluorimeter and RNA integrity number (RIN) on an Agilent TapeStation. RNA was stored at -80°C and transported to the genomic facility, Centre for Genome Enabled Biology and Medicine, at University of Aberdeen on dry ice. cDNA libraries were prepared with 50ng input at 14 cycles of amplification using Illumina Stranded mRNA Prep, Ligation kit (20040532) and sequenced on a NextSeq 500 machine.
FASTQ files of single-end reads were processed using FASTP (*17*). Processed reads were mapped to human reference genome assembly GRCh38 and transcript level quantification was obtained using Salmon v1.10.2 (*18*). Transcript-level quantification data was imported in R, and using tximeta (*19*) Bioconductor package (https://bioconductor.org/packages/3.19/bioc/html/tximeta.html) a *SummarizedExperiment* (SE) containing gene-level quantification, experimental metadata and annotation metadata for genes was obtained. SummarizedExperiment was loaded into the DESeq2 package (*20*) using the DESeqDataSet function. For DESeq2 analysis, experimental design included experimental group and donor. Genes with at least two non-zero counts across samples were retained. DESeq2 differential expression pipeline was run by calling function DESeq and result tables containing gene symbol, ensemble gene ID, base mean, log2 fold change (log2FC), p value, and false discovery rate (FDR)- adjusted p value (adj P value) were exported in csv files. Volcano plots were generated using ggplot2 (version 3.5.1) and ggrepel (version 0.9.6) packages.

**Protein fractionation and proteomic analysis**

For proteomic analysis using LC-MS/MS, vehicle- and IL-17A-treated day 8 cerebroids were collected from three donors and snap frozen in liquid nitrogen. Protein extraction and fractionation **(Fig. 4G)** was performed using a previously published three-step decellularization and ECM extraction protocol that yields cellular (CHAPS), soluble ECM (sECM) and insoluble ECM (iECM) fractions (*21*).

Protein extracts were processed using the filter-aided sample preparation (FASP) Protein Digestion Kit (Abcam, Cat ab270519) according to the manufacturer's instructions with the following modifications. Step 6.1 was repeated until the whole sample was loaded onto the filter. In Step 6.9, 50 µl of 20 µg/ml Sequencing Grade Trypsin (Promega Cat V5111) in 50 mM ammonium bicarbonate was added to 25 µl of 50mM ammonium bicarbonate on the spin filter. Each sample was therefore treated with 1 µg trypsin, optimised for a total protein content of 30 µg.

Following FASP digestion and collection of the peptides, the samples were snap frozen at -70 °C and dried in a vacuum centrifuge (SpeedVac). Samples were dissolved in 0.5% TFA and desalted using Merck ZipTip Pipette Tips (Cat ZTC18M960). The eluted peptides were dried in a SpeedVac and resuspended in 10 µl 0.1% TFA.

The Liquid chromatography – mass spectrometry (LC-MS) system comprised an UltiMate 3000 RSLCnano coupled to a Q Exactive Plus (Thermo Fisher Scientific). Peptides (5 µl) were injected via a sample loop using loading pump buffer A (0.1% TFA in ultrapure water) at 10 µl/min and bound to the C18 PepMap 100 precolumn (300 µm x 5 mm, Thermo Scientific P/N 160454). After 5 min, flow was switched to the nano pump and peptides were reverse flushed to the PepMap RSLC C18 column (75 µm x 25 cm, Thermo Scientific P/N ES801) and separated with a gradient of acetonitrile increasing from 2.4% to 32% in 35 min. Nano pump solvent A was 0.1% formic acid in ultrapure water while nano pump solvent B was 0.1% formic acid in 80% acetonitrile. The column was washed for 8 minutes in 90% solvent B and then equilibrated for 15 minutes in 3% solvent B prior to the next injection.

The Q Exactive Plus was operated in positive polarity mode using a Top 10 data-dependent acquisition (DDA) method. Full MS scan parameters were Resolution = 70,000; AGC Target = 3e6; Maximum IT = 50 ms; Scan Range = 375-1750 m/z. dd-MS2 parameters were: Resolution = 17,500; AGC Target = 5e4; Maximum IT = 100 ms; Loop Count = 10; Isolation Window = 1.6 m/z; NCE = 28; Charge Exclusion = <2 and >5; Peptide Match = preferred; Dynamic Exclusion = 40 s.

MS raw data files, acquired between 5-65 min after flow switching, were processed using Proteome Discoverer (version 2.2, Thermo Scientific) using a workflow that incorporated Mascot Server (version 2.6, Matrix Science), target decoy peptide-spectrum match validation (strict false discovery rate of 0.01) and peptide quantification by precursor ion peak area. Protein-level abundance quantification data was imported in R. For each fraction, proteins with non-zero abundance in at least two samples were included in the analysis. Protein abundance values were log-transformed and normalized with median centered method of data normalization. Missing values were imputed by random draws from a Gaussian distribution centered to a minimal value using impute.MinProb() command in imputeLCMD package v2.1. SummarizedExperiment package was used to generate SE object and limma (*22*) package (version 3.19) was used to identify differentially abundant proteins by applying a multi-factorial design assay including the treatment group and donor. Volcano plots were generated using ggplot2 and ggrepel packages.

**Meta-analysis of RNA seq data**

We searched PubMed and Web of Science for studies published in English from database inception to May 30, 2023, that investigated the transcriptome of human ASD brains. We used the search terms ("ASD" OR "autism") AND ("brain" OR "cortex") AND ("transcriptome" OR "RNAseq" OR "microarray" OR "gene expression"). This led to identification of 3 microarray (*23-25*), 6 bulk (*26-31*), and 3 single nuclear RNA sequencing (*30, 32, 33*) studies on the post-mortem brain tissue of the ASD cases and controls. Gandal et al (*29*) performed a meta-analysis of some published studies (*23-25*) and identified a list of DEGs. Data from one study (*28*), after applying the filters, gave a list of only 6 upregulated genes (2 mitochondrial genes and 4 small nucleolar RNA genes) and was excluded from further analysis. Studies that included bulk RNA-seq data from the frontal cortex (*27, 29, 30*) and meta-analysis of microarray data (*29*) were selected for further analysis. To identify functionally enriched terms and pathways in an unbiased manner, we applied a protein sub-network based functional enrichment analysis using STRING v12.0 (*34*). Functional enrichment analysis was performed for each individual study with genes ranked according to log2FC in ASD versus control brains. STRING tests the protein of each known pathway for any non-random skew within the provided input and thus tests pathways for a skewed distribution on either end of this ranked input. It applies Kolmogorov–Smirnov testing to detect statistically significant pathway s(*34, 35*). Terms enriched in at least two independent studies, with 50 or more genes, and an FDR-adjusted p value < 0.05 were considered significant and plotted **(fig. S13)**.

**Functional enrichment analysis**

For transcriptomic and proteomic data, we applied a protein sub-network based functional enrichment analysis using STRING v12.0 as described above. Terms enriched in Reactome database are plotted **(Fig. 4F and fig. S8-11)**. K-means clustering was performed on protein-protein interaction network of differentially abundant proteins using STRING.
Over-representation analysis (ORA) was performed using Enrichr (*36*) (https://maayanlab.cloud/Enrichr/). For ORA, genes with log2FC above 0.4 and below -0.4 and p value below 0.05 were considered up- and down-regulated, respectively. All genes detected by RNA-seq were provided as background genes. Terms with FDR-adjusted p value below 0.05 were considered significant. Enrichr was used to identify list of enriched transcription factors (TFs) having regulatory interactions with DEGs in human- TRRUST v2 (Transcriptional Regulatory Relationships Unraveled by Sentence-based Text mining) reference database (*37*). Data visualization was performed in R using ggplot2.
The list of genes implicated in neurological disorders is adopted from a recent pre-print (*38*).

**Statistical analysis and data visualization**

GraphPad Prism (version 8.4.3) and R (version 4.3.1) statistical computing environment (https://www.r-project.org/) were used to analyze data and generate graphs. Throughout the paper, * indicates p < 0.05, ** indicates p < 0.01, *** indicates p < 0.005, **** indicates p < 0.001. Two-way Analysis of Variance (ANOVA) test was applied to compare across experimental groups (vehicle- versus IL-17A-treated) and donors. For inter-group comparison across multiple groups, Tukey's multiple comparisons test was performed. Donor-level quantification is represented as mean ± Standard Error of Mean (SEM) while individual data points represent technical replicates. On average, 2-3 sections per cerebroids and 2-3 cerebroids per donor per group were analyzed for each quantification. The number of donors for each quantification are provided in the figure legends.
Plots were made in GraphPad Prism or R and edited in Inkscape v 1.3.2.

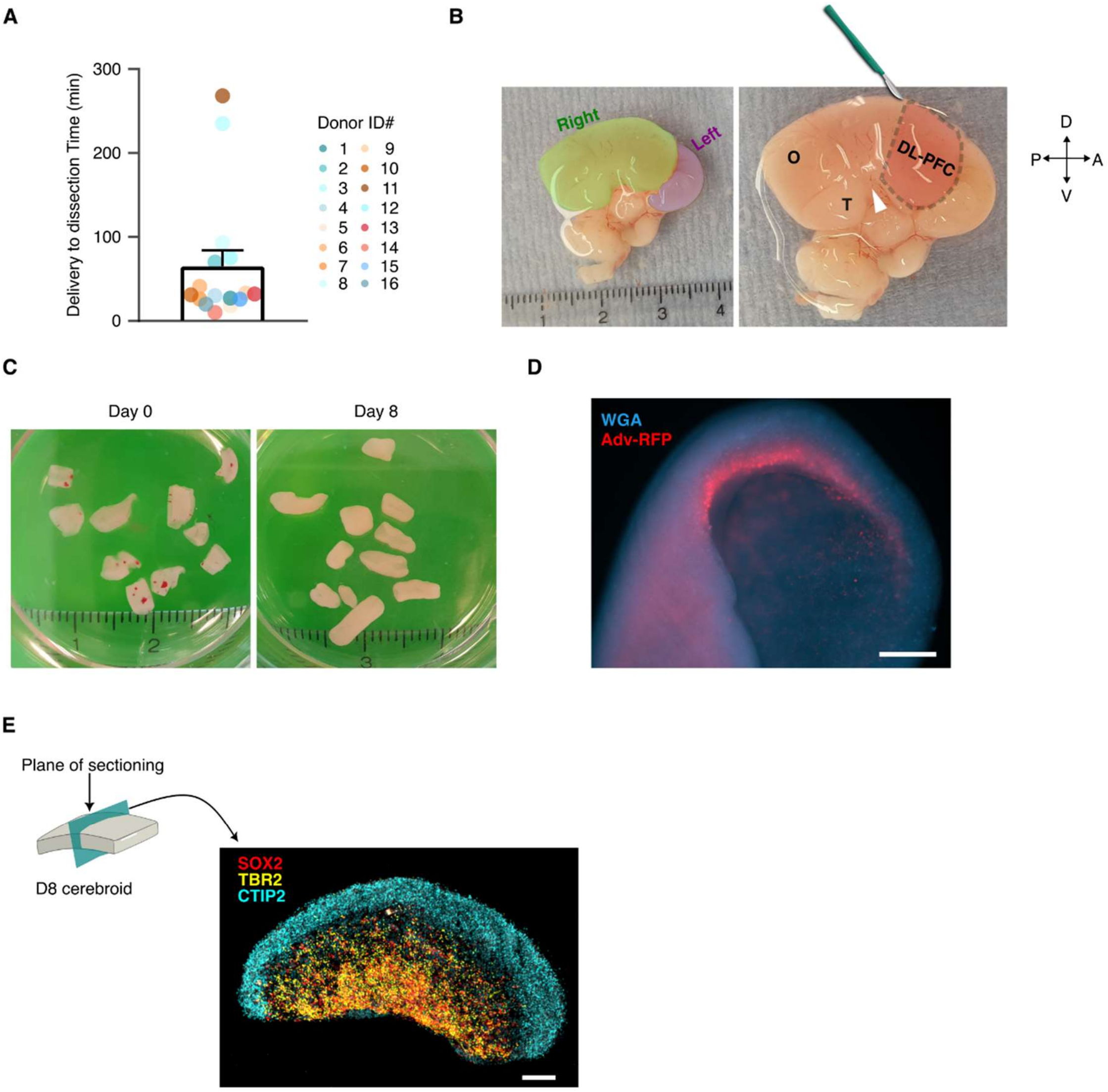


**Fig.S1. Generation of cerebroids from human dorsolateral prefrontal cortex (DLPFC) tissue.**

**(A)** Time between delivery of the fetus and brain dissection recorded for 14 donors (in minutes). Data is represented as mean ± SEM and each data point represents individual donor. (See table S1 for donor details).

**(B)** Dissected human brain at gestational week (GW) 12. The right hemisphere is highlighted in green and the left hemisphere in purple, with the DL-PFC outlined by a dashed line. The occipital (O) and temporal (T) lobes are labeled, and the lateral sulcus is marked by an arrowhead. Brain orientation: posterior (P), anterior (A), dorsal (D), and ventral (V). A ruler (in

cm) is shown for scale. For cerebroid culture, DLPFC was identified by the anatomical landmarks shown here and dissected.
**(C)** Images of cerebroids immediately after dissection (Day 0) and following 8 days in culture (Day 8). Note the increase in the size of the cerebroids after 8 days in culture.
**(D)** Fluorescence image of Day 8 cerebroids stained with Wheat Germ Agglutinin (WGA), marking glycocalyx and proteoglycans in the extracellular matrix (ECM), and adenovirus-RFP-labeled neural stem cells (NSCs) lining the ventricular surface. Scale bar, 500μm.
**(E)** Schematic of the sectioning plane and a fluorescence image of a whole cerebroid section stained for SOX2 (NSCs), TBR2 (intermediate progenitor cells, IPCs), and CTIP2 (neurons).

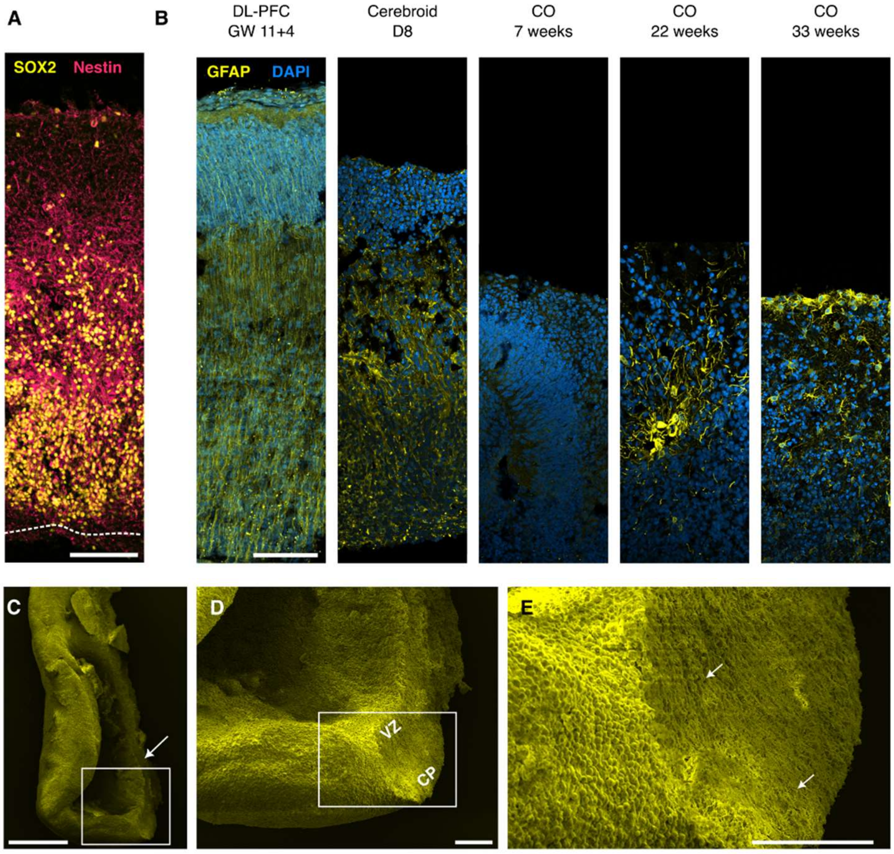


**Fig. S2. Characterization of NSCs in cerebroid cultures.**
**(A)** Representative fluorescence image of $SOX2^+$ and $Nestin^+$ neural stem cells (NSCs) in cerebroid culture. Scale bar, 100 µm.
**(B)** Fluorescence images showing GFAP-expressing radial glia (RG) in the human dorsolateral prefrontal cortex (DL-PFC), day 8 (D8) cerebroids, and cortical organoids (COs) cultured for 7, 22, and 33 weeks. Note the absence of GFAP staining in the NSC layer of 7-week-old COs, while only $GFAP^+$ parenchymal astrocytes are observed in 22- and 33-week-old COs. This contrasts with $GFAP^+$ processes of NSCs in DL-PFC and D8 cerebroids. Scale bar, 100 µm.
**(C–E)** Scanning electron microscopy (SEM) images of day 8 cerebroids. (C) Low-magnification image showing the cut surface of a cerebroid. (D) Higher-magnification image highlighting cytoarchitectural features, including the ventricular zone (VZ) and cortical plate (CP). (E) SEM image showing radial glial cell processes (white arrows). Scale bars: 500 µm (C), 100 µm [(D), (E)].

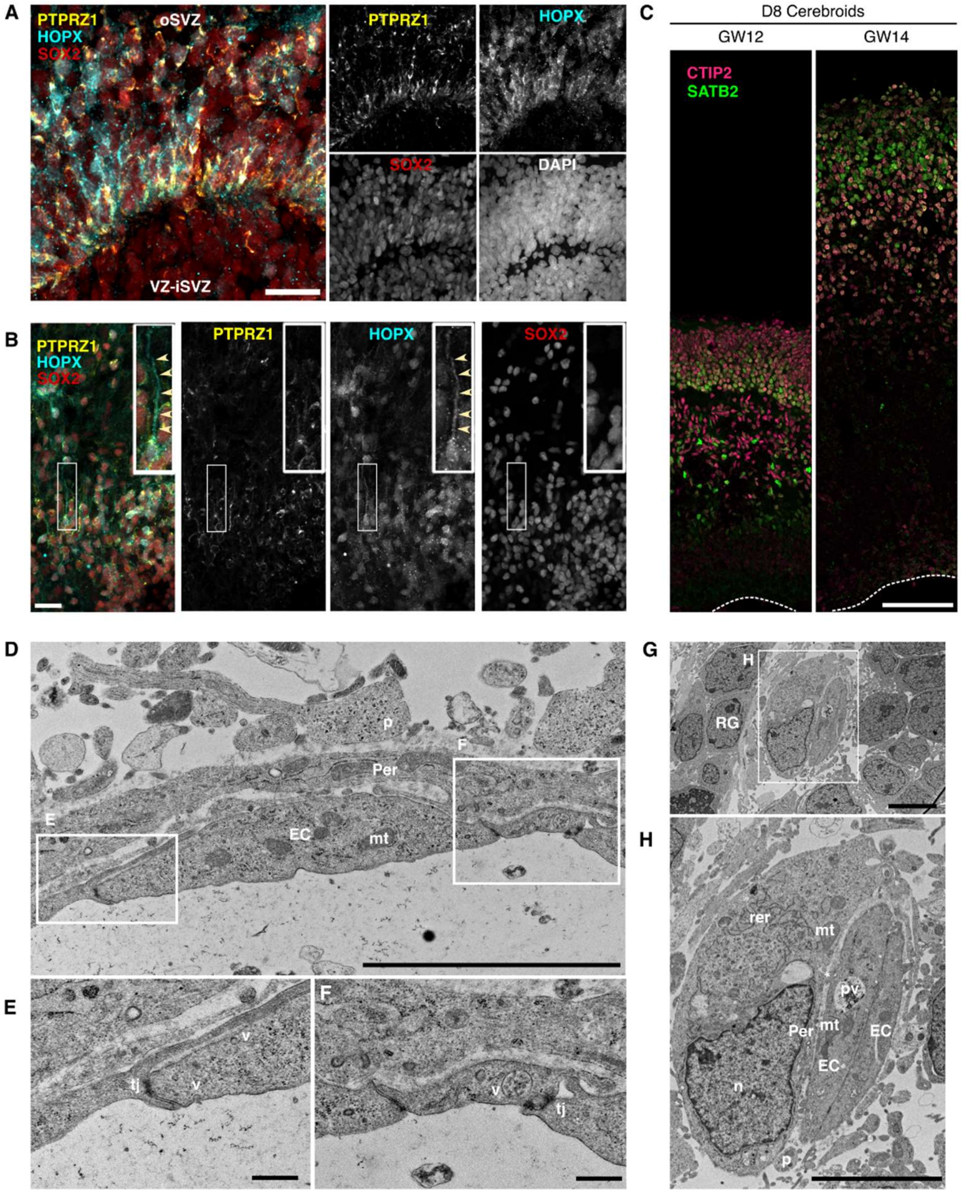


**Fig. S3. Characterization of NSCs and vascular structures in cerebroid cultures.**
**(A and B)** Representative fluorescence images showing outer radial glial cells (oRGs) in the outer subventricular zone (oSVZ) of cerebroids cultured for 8 days. Ventricular zone (VZ) and inner subventricular zone (iSVZ) are indicated. oRGs express the markers PTPRZ1 and HOPX. Yellow arrowheads highlight the $HOPX^{+}$ processes of oRGs. Scale bars: 100 µm (A) and 25 µm (B).
**(C)** Fluorescence images showing the expression of neuronal markers in Day 8 cerebroids from 12- and 14-week-old fetuses. Neuronal markers include CTIP2 (deep-layer neurons), and SATB2 (late-born upper-layer neurons). Scale bar, 100 µm.
**(D–F)** TEM images of vascular structures in Day 2 cerebroids. (D) Annotated image from Fig.

1U showing an endothelial cell (EC) with mitochondria (mt), surrounded by a pericyte (Per) and the end-foot process of a radial glial cell (RGC, p). Zoomed-in views show tight junctions (tj) between endothelial cells, forming part of the blood-brain barrier, and secretory vesicles (v). Scale bar, 5 µm (D), and 500 nm [(E), (F)].

**(G and H)** TEM images of a vascular unit lacking a lumen, showing a radial glial cell (RG), endothelial cell (EC), and pericyte (Per). Both endothelial cells and pericytes are rich in mitochondria (mt). The endothelial cell contains a phagocytic vesicle (pv), while the pericyte shows a high content of rough endoplasmic reticulum (rer). A peg-socket junction (*) between the pericyte and endothelial cell is also visible. Scale bar, 5 µm.

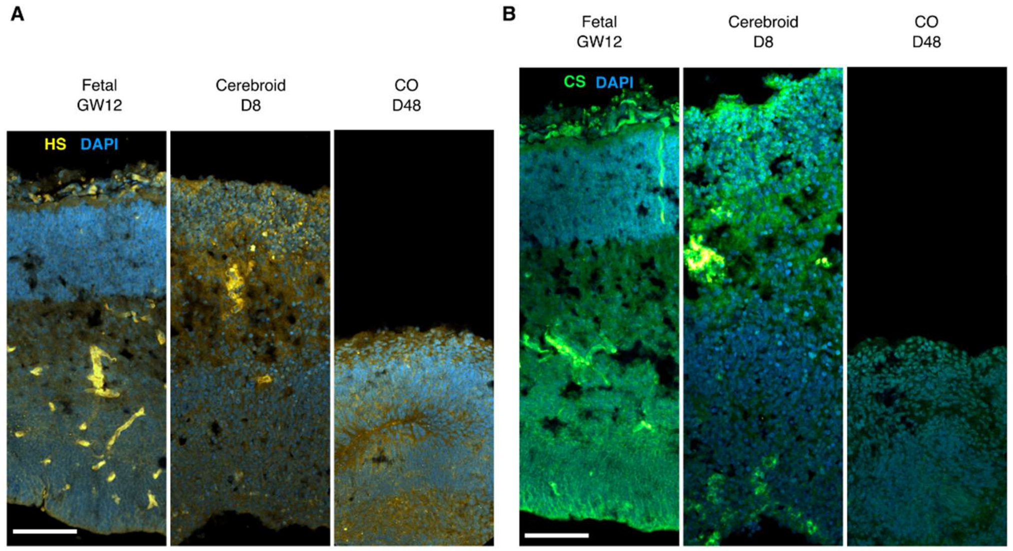


**Fig. S4. Comparison of ECM components in fetal, cerebroid, and cortical organoid tissues.**
**(A)** Representative fluorescence images of Heparan Sulfate (HS) staining in a GW12 dorsolateral prefrontal cortex (DL-PFC) section, an 8-day cultured cerebroid, and a 48-day-old cortical organoid (CO). HS fluorescence intensities are higher and more abundant in DL-PFC and cerebroids as compared to CO.
**(B)** Representative fluorescence images of Chondroitin Sulfate (CS) staining in a GW12 DL-PFC section, an 8-day cultured cerebroid, and a 48-day-old CO. In contrast to CO, cerebroids show greater abundance of CS mimicking expression pattern in DL-PFC.
Scale bar, 100 µm

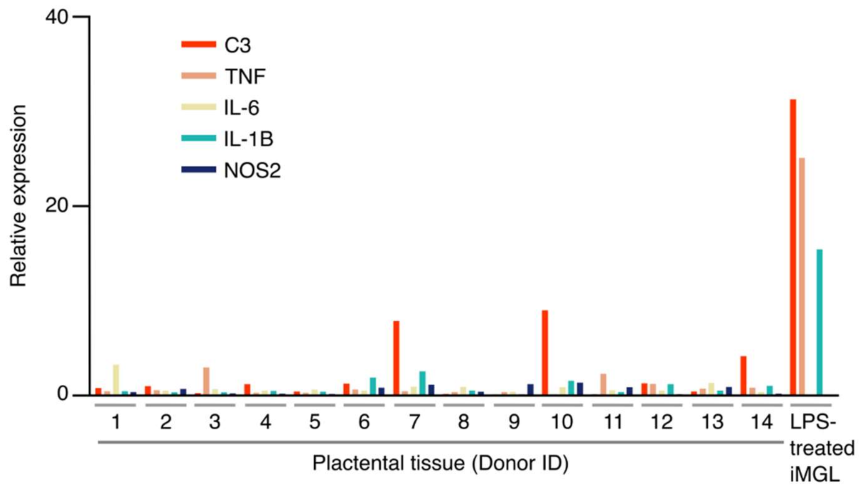


**Fig. S5. qPCR analysis of inflammatory markers in donor placental tissue.**
qPCR analysis of inflammatory marker expression in placental tissue from respective donors, compared to LPS-treated iPSC-derived microglia like cells (iMGL).

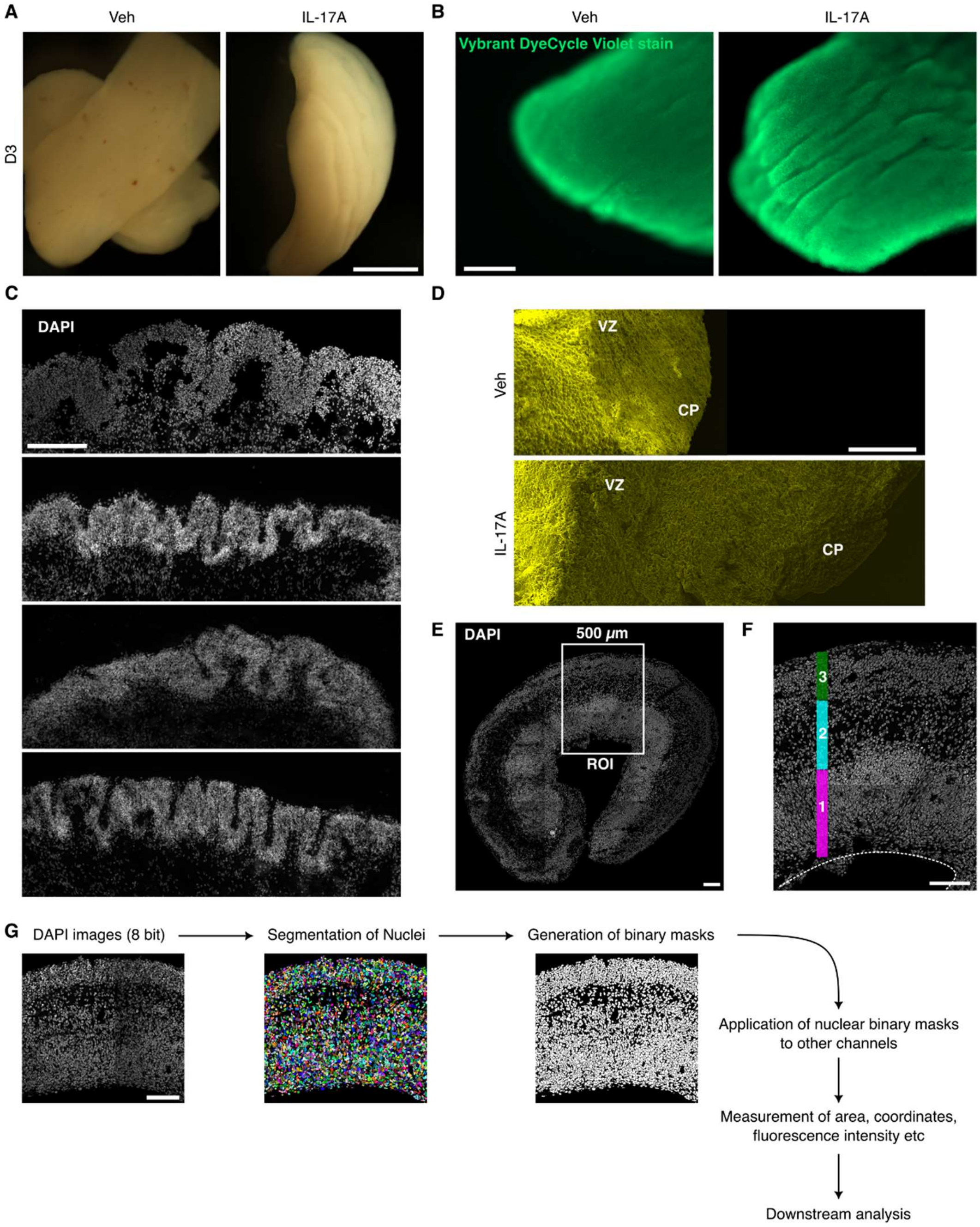


**Fig. S6. Characterization of IL-17A-induced phenotypes in cerebroids.**
**(A)** Representative images of vehicle- and IL-17A-treated cerebroids after 3 days of culture. Cortical folding is evident in IL-17A-treated day 3 cerebroid. Scale bar, 1000 µm.
**(B)** Representative live fluorescence images of vehicle- and IL-17A-treated day 8 cerebroids stained with a nuclear dye (Vybrant DyeCycle Violet). IL-17A-treated cerebroids show abundant and deep folding on the cortical surface in contrast to occasional, shallow folds in vehicle-treated

cerebroids. Scale bar, 500 μm.
**(C)** Examples of various gyrification patterns in Day 8 IL-17A-treated cerebroids, visualized by DAPI staining. Scale bar, 100 μm.
**(D)** SEM images of cut surfaces of Day 8 vehicle- and IL-17A-treated cerebroids, showing increased thickness in IL-17A-treated cerebroids. Scale bar, 100 μm.
**(E and F)** Image-based quantifications for cell numbers and fluorescence intensities. Representative images of DAPI-stained Day 8 cerebroid sections are shown. (E) A fixed-width region of interest (ROI) was drawn through the central part of the cerebroid for image-based quantifications. Scale bar, 100 μm. (F) Magnified view of the ROI from (E). The dotted line represents the ventricular surface. Three layers were identified based on cell densities and immunofluorescence markers: SOX2-rich layer 1 and CTIP2-rich layer 3 (as shown in Fig. 1B). Scale bar, 100 μm.
**(G)** Schematic of the image analysis pipeline for fluorescence intensity-based analysis (as shown in Fig. 2M and N). 8-bit images of DAPI-stained nuclei were used to segment individual nuclei, followed by the generation of binary masks. These masks were then split to generate ROIs for individual nuclei, and fluorescence intensities were measured across these nuclear ROIs for various channels.

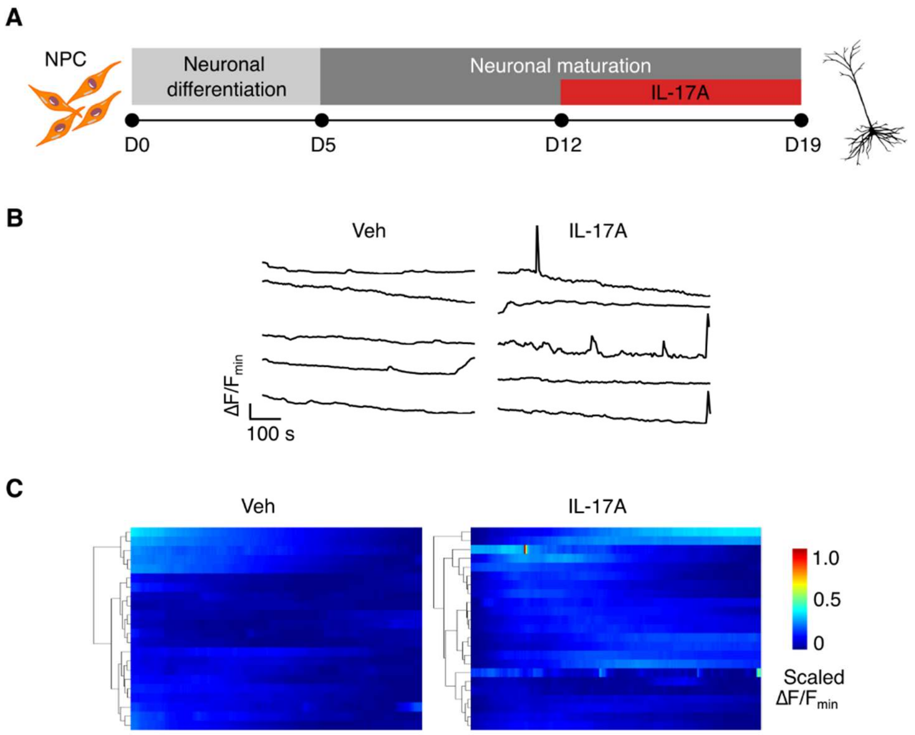


**Fig. S7. IL-17A-treatment increases calcium transients in 2D cultured iPSC-derived neurons**

**(A)** Schematic of the experimental paradigm for 2D neuronal culture and IL-17A treatment. iPSC-derived neural progenitor cells (NPCs) were differentiated into forebrain neurons, followed by neuronal maturation. From Day 12 to Day 19, neurons were treated with 10 ng/mL IL-17A every 2 days to assess the effects of IL-17A on neuronal maturity.

**(B)** Representative calcium transient plots in vehicle- and IL-17A-treated neurons. Each tracing represents the ratio of change in mean fluorescence intensity (MFI) from the minimum MFI ($\Delta F$) to the minimum MFI ($F_{min}$) over time. IL-17A-treated neurons exhibit increased amplitude and frequency of spontaneous calcium transient activity compared to vehicle-treated controls.

**(C)** Heatmaps of calcium transients in vehicle- and IL-17A-treated neurons. Each row represents an individual neuron, while each column represents the time scale. Colors indicate the scaled $\Delta F/F_{min}$ values over time, highlighting the heightened calcium activity in IL-17A-treated neurons.

A

B

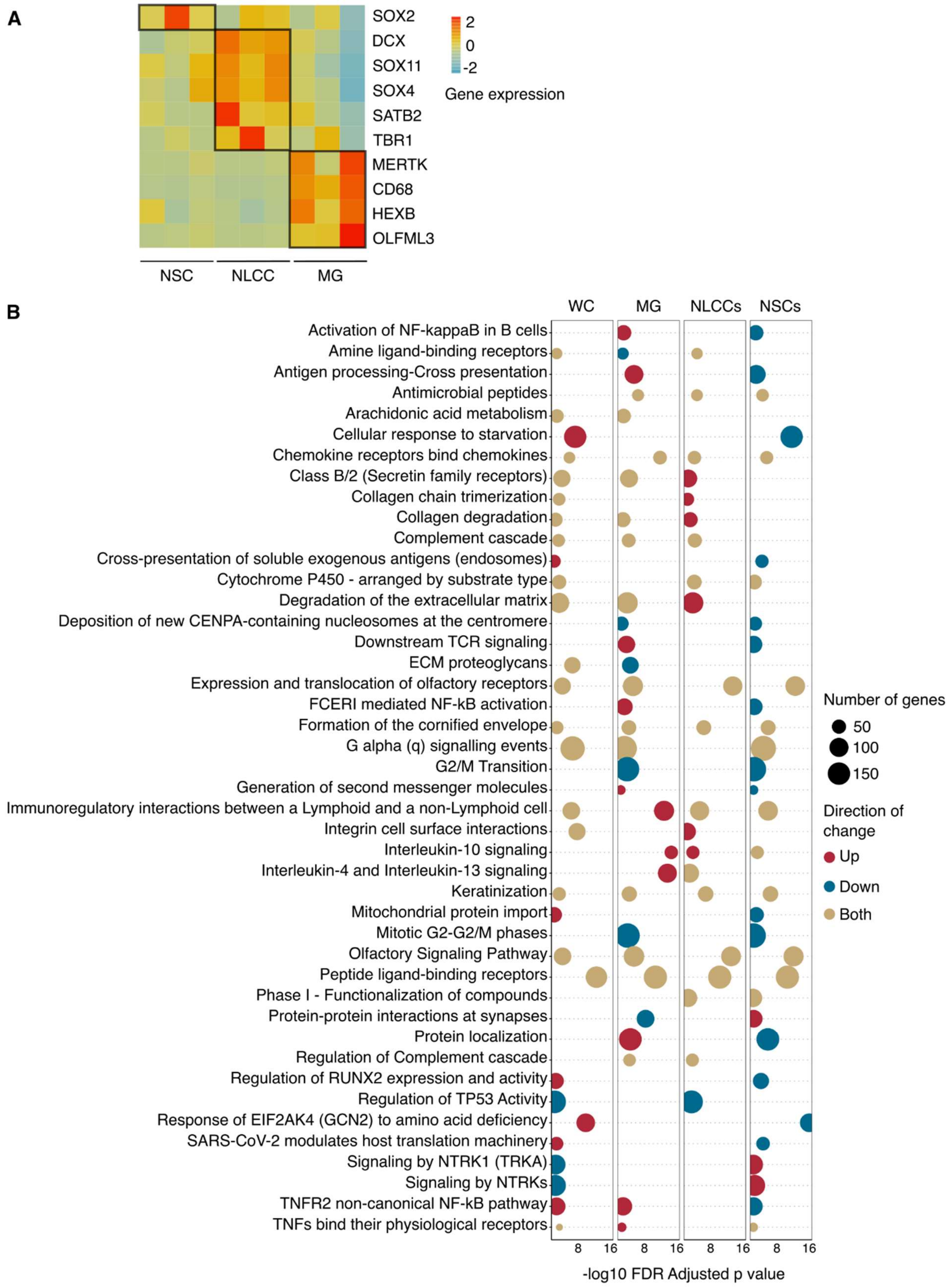
SOX2
DCX
SOX11
SOX4
SATB2
TBR1
MERTK
CD68
HEXB
OLFML3
2
0
-2
Gene expression
NSC
NLCC
MG
WC
MG
NLCCs
NSCs
Activation of NF-kappaB in B cells
Amine ligand-binding receptors
Antigen processing-Cross presentation
Antimicrobial peptides
Arachidonic acid metabolism
Cellular response to starvation
Chemokine receptors bind chemokines
Class B/2 (Secretin family receptors)
Collagen chain trimerization
Collagen degradation
Complement cascade
Cross-presentation of soluble exogenous antigens (endosomes)
Cytochrome P450 - arranged by substrate type
Degradation of the extracellular matrix
Deposition of new CENPA-containing nucleosomes at the centromere
Downstream TCR signaling
ECM proteoglycans
Expression and translocation of olfactory receptors
FCERI mediated NF-kB activation
Formation of the cornified envelope
G alpha (q) signalling events
G2/M Transition
Generation of second messenger molecules
Immunoregulatory interactions between a Lymphoid and a non-Lymphoid cell
Integrin cell surface interactions
Interleukin-10 signaling
Interleukin-4 and Interleukin-13 signaling
Keratinization
Mitochondrial protein import
Mitotic G2-G2/M phases
Olfactory Signaling Pathway
Peptide ligand-binding receptors
Phase I - Functionalization of compounds
Protein-protein interactions at synapses
Protein localization
Regulation of Complement cascade
Regulation of RUNX2 expression and activity
Regulation of TP53 Activity
Response of EIF2AK4 (GCN2) to amino acid deficiency
SARS-CoV-2 modulates host translation machinery
Signaling by NTRK1 (TRKA)
Signaling by NTRKs
TNFR2 non-canonical NF-kB pathway
TNFs bind their physiological receptors
8 16 8 16 8 16 8 16
-log10 FDR Adjusted p value
Number of genes
50
100
150
Direction of change
Up
Down
Both

**Fig. S8. Marker genes for sorted cell populations and functional enrichment analysis of transcriptomic data.**

**(A)** Heatmap showing scaled expression of selected marker genes in neural stem cells (NSCs, SOX2), neuron-lineage committed cells (NLCCs, DCX, SOX11, SOX4, SATB2, TBR1), and microglia (MG, MERTK, CD68, HEXB, OLFML3) isolated from Day 8 cerebroids (n = 3 donors). The heatmap highlights the distinct expression profiles of these marker genes across the sorted cell populations.

**(B)** Bubble plot illustrating enriched Reactome pathways identified in two or more RNA-seq datasets (whole cerebroid, MG, NLCCs, and NSCs). Pathway enrichment was determined using protein network-based gene set enrichment analysis via STRING, with genes ranked by Log2FC. Only pathways enriched in at least two datasets and involving ≥50 genes are shown. Each column represents a transcriptomic dataset, with the position of the bubble along the x-axis indicating the -log10 of the FDR-adjusted p-value. Bubble size corresponds to the number of genes associated with each pathway, while color denotes the direction of gene expression changes

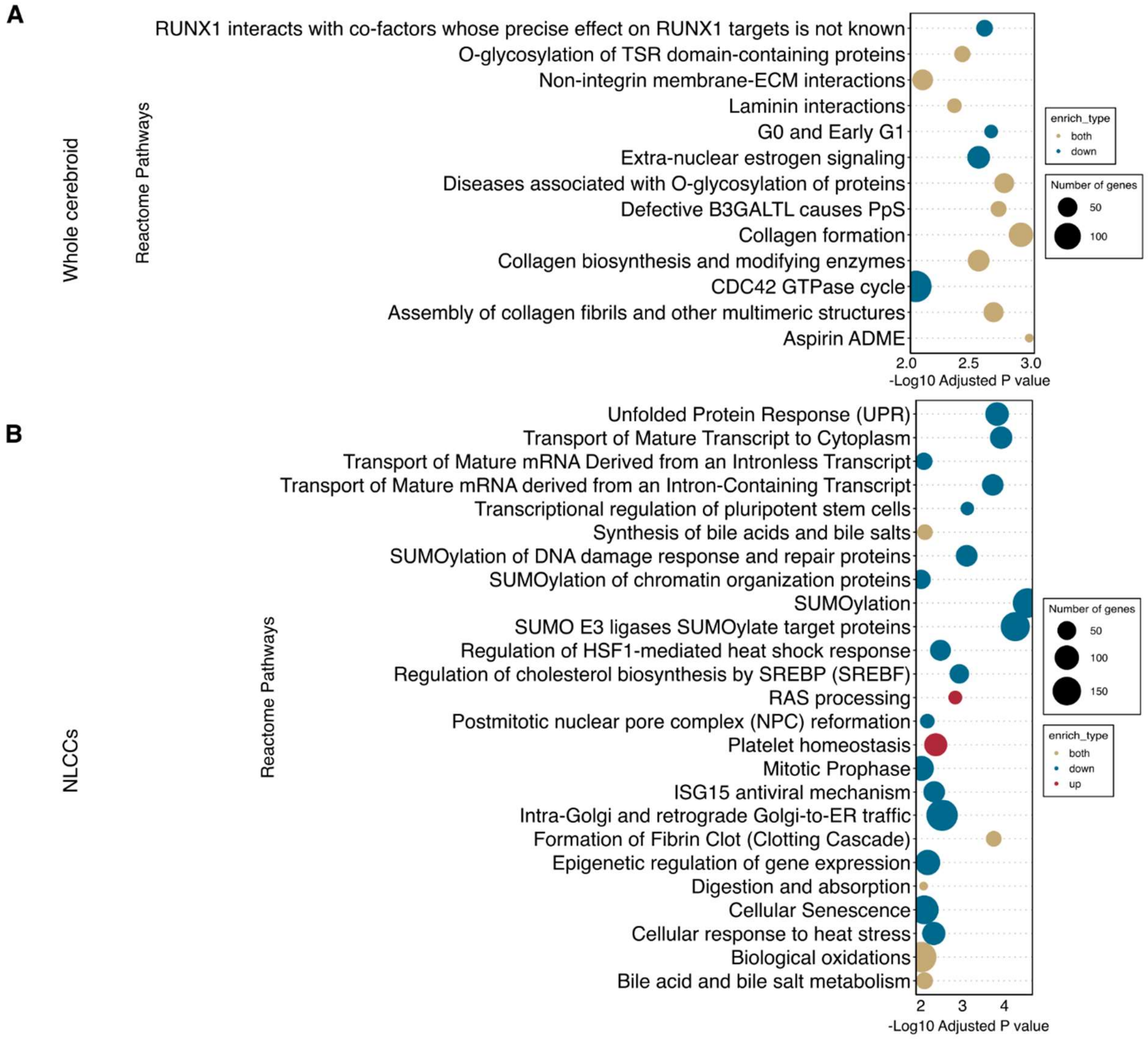


**Fig. S9. Functional enrichment analysis of whole cerebroid and neuronal transcriptomic data.**

(**A and B**) Uniquely enriched Reactome pathways in whole cerebroid tissue (A) and neuron-lineage committed cells (NLCCs) (B). Pathway enrichment was determined using protein network-based gene set enrichment analysis via STRING, with genes ranked by Log2FC. The position of the bubble along the x-axis indicating the -log10 of the FDR-adjusted p-value. Bubble size corresponds to the number of genes associated with each pathway, while color denotes the direction of gene expression changes.

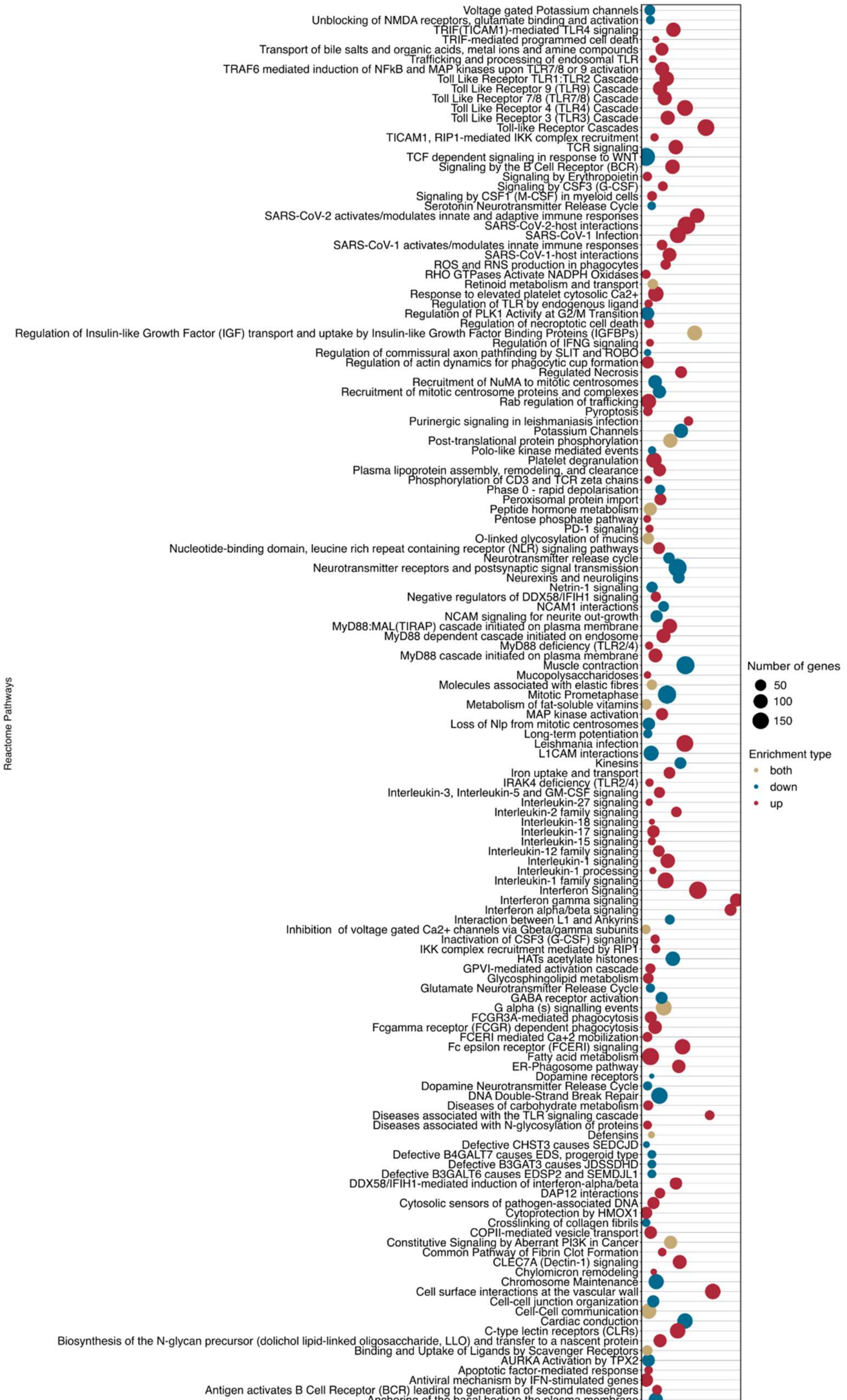
Voltage gated Potassium channels
Unblocking of NMDA receptors, glutamate binding and activation
TRIF(TICAM1)-mediated TLR4 signaling
TRIF-mediated programmed cell death
Transport of bile salts and organic acids, metal ions and amine compounds
Trafficking and processing of endosomal TLR
TRAF6 mediated induction of NFkB and MAP kinases upon TLR7/8 or 9 activation
Toll Like Receptor TLR1:TLR2 Cascade
Toll Like Receptor 9 (TLR9) Cascade
Toll Like Receptor 7/8 (TLR7/8) Cascade
Toll Like Receptor 4 (TLR4) Cascade
Toll Like Receptor 3 (TLR3) Cascade
Toll-like Receptor Cascades
TICAM1, RIP1-mediated IKK complex recruitment
TCR signaling
TCF dependent signaling in response to WNT
Signaling by the B Cell Receptor (BCR)
Signaling by Erythropoietin
Signaling by CSF3 (G-CSF)
Signaling by CSF1 (M-CSF) in myeloid cells
Serotonin Neurotransmitter Release Cycle
SARS-CoV-2 activates/modulates innate and adaptive immune responses
SARS-CoV-2-host interactions
SARS-CoV-1 Infection
SARS-CoV-1 activates/modulates innate immune responses
SARS-CoV-1-host interactions
ROS and RNS production in phagocytes
RHO GTPases Activate NADPH Oxidases
Retinoid metabolism and transport
Response to elevated platelet cytosolic Ca2+
Regulation of TLR by endogenous ligand
Regulation of PLK1 Activity at G2/M Transition
Regulation of necroptotic cell death
Regulation of Insulin-like Growth Factor (IGF) transport and uptake by Insulin-like Growth Factor Binding Proteins (IGFBPs)
Regulation of IFNG signaling
Regulation of commissural axon pathfinding by SLIT and ROBO
Regulation of actin dynamics for phagocytic cup formation
Regulated Necrosis
Recruitment of NuMA to mitotic centrosomes
Recruitment of mitotic centrosome proteins and complexes
Rab regulation of trafficking
Pyroptosis
Purinergic signaling in leishmaniasis infection
Potassium Channels
Post-translational protein phosphorylation
Polo-like kinase mediated events
Platelet degranulation
Plasma lipoprotein assembly, remodeling, and clearance
Phosphorylation of CD3 and TCR zeta chains
Phase 0 - rapid depolarisation
Peroxisomal protein import
Peptide hormone metabolism
Pentose phosphate pathway
PD-1 signaling
O-linked glycosylation of mucins
Nucleotide-binding domain, leucine rich repeat containing receptor (NLR) signaling pathways
Neurotransmitter release cycle
Neurotransmitter receptors and postsynaptic signal transmission
Neurexins and neuroligins
Netrin-1 signaling
Negative regulators of DDX58/IFIH1 signaling
NCAM1 interactions
NCAM signaling for neurite out-growth
MyD88:MAL(TIRAP) cascade initiated on plasma membrane
MyD88 dependent cascade initiated on endosome
MyD88 deficiency (TLR2/4)
MyD88 cascade initiated on plasma membrane
Muscle contraction
Mucopolysaccharidoses
Molecules associated with elastic fibres
Mitotic Prometaphase
Metabolism of fat-soluble vitamins
MAP kinase activation
Loss of Nlp from mitotic centrosomes
Long-term potentiation
Leishmania infection
L1CAM interactions
Kinesins
Iron uptake and transport
IRAK4 deficiency (TLR2/4)
Interleukin-3, Interleukin-5 and GM-CSF signaling
Interleukin-27 signaling
Interleukin-2 family signaling
Interleukin-18 signaling
Interleukin-17 signaling
Interleukin-15 signaling
Interleukin-12 family signaling
Interleukin-1 signaling
Interleukin-1 processing
Interleukin-1 family signaling
Interferon Signaling
Interferon gamma signaling
Interferon alpha/beta signaling
Interaction between L1 and Ankyrins
Inhibition of voltage gated Ca2+ channels via Gbeta/gamma subunits
Inactivation of CSF3 (G-CSF) signaling
IKK complex recruitment mediated by RIP1
HATs acetylate histones
GPVI-mediated activation cascade
Glycosphingolipid metabolism
Glutamate Neurotransmitter Release Cycle
GABA receptor activation
G alpha (s) signalling events
FCGR3A-mediated phagocytosis
Fcgamma receptor (FCGR) dependent phagocytosis
FCERI mediated Ca+2 mobilization
Fc epsilon receptor (FCERI) signaling
Fatty acid metabolism
ER-Phagosome pathway
Dopamine receptors
Dopamine Neurotransmitter Release Cycle
DNA Double-Strand Break Repair
Diseases of carbohydrate metabolism
Diseases associated with the TLR signaling cascade
Diseases associated with N-glycosylation of proteins
Defensins
Defective CHST3 causes SEDCJD
Defective B4GALT7 causes EDS, progeroid type
Defective B3GAT3 causes JDSSDHD
Defective B3GALT6 causes EDSP2 and SEMDJL1
DDX58/IFIH1-mediated induction of interferon-alpha/beta
DAP12 interactions
Cytosolic sensors of pathogen-associated DNA
Cytoprotection by HMOX1
Crosslinking of collagen fibrils
COPII-mediated vesicle transport
Constitutive Signaling by Aberrant PI3K in Cancer
Common Pathway of Fibrin Clot Formation
CLEC7A (Dectin-1) signaling
Chylomicron remodeling
Chromosome Maintenance
Cell surface interactions at the vascular wall
Cell-cell junction organization
Cell-Cell communication
Cardiac conduction
C-type lectin receptors (CLRs)
Biosynthesis of the N-glycan precursor (dolichol lipid-linked oligosaccharide, LLO) and transfer to a nascent protein
Binding and Uptake of Ligands by Scavenger Receptors
AURKA Activation by TPX2
Apoptotic factor-mediated response
Antiviral mechanism by IFN-stimulated genes
Antigen activates B Cell Receptor (BCR) leading to generation of second messengers
Anchoring of the basal body to the plasma membrane
Adherens junctions interactions
Activation of Matrix Metalloproteinases
Activation of G protein gated Potassium channels
A tetrasaccharide linker sequence is required for GAG synthesis
Reactome Pathways
2 4 6 8 10
-Log10 Adjusted P value
Number of genes
50
100
150
Enrichment type
both
down
up

**Fig. S10. Unique enriched Reactome pathways in microglia from IL-17A-treated cerebroids.**

Uniquely enriched Reactome pathways for microglia isolated from IL-17A-treated cerebroids. Pathway enrichment was determined using protein network-based gene set enrichment analysis via STRING, with genes ranked by Log2FC. The position of the bubble along the x-axis indicating the -log10 of the FDR-adjusted p-value. Bubble size corresponds to the number of genes associated with each pathway, while color denotes the direction of gene expression changes.

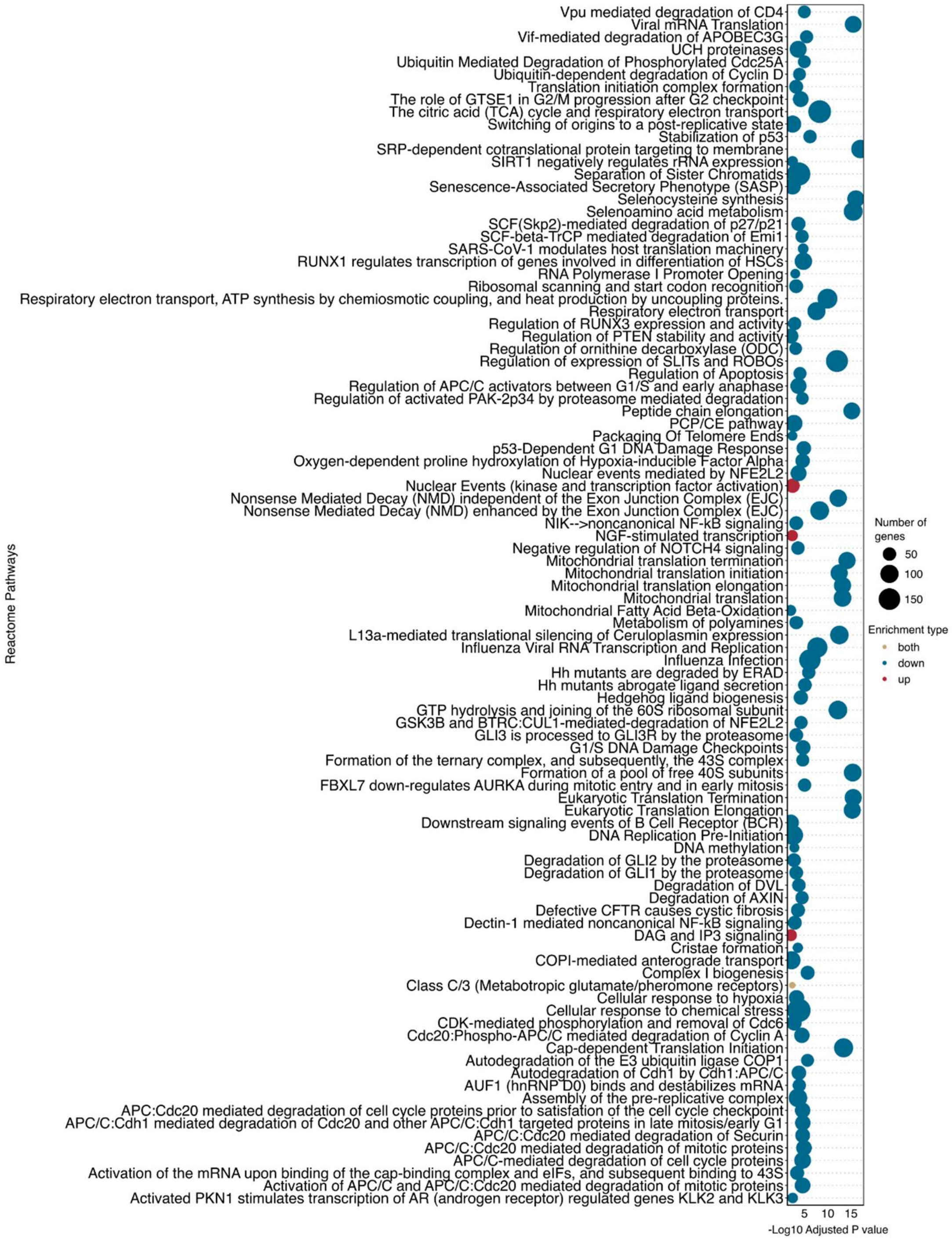


**Fig. S11. Unique enriched Reactome pathways in neural stem cells from IL-17A-treated cerebroids.**

Uniquely enriched Reactome pathways for neural stem cells (NSCs) isolated from IL-17A-treated cerebroids. Pathway enrichment was determined using protein network-based gene set enrichment analysis via STRING, with genes ranked by Log2FC. The position of the bubble along the x-axis indicating the -log10 of the FDR-adjusted p-value. Bubble size corresponds to the number of genes associated with each pathway, while color denotes the direction of gene expression changes.

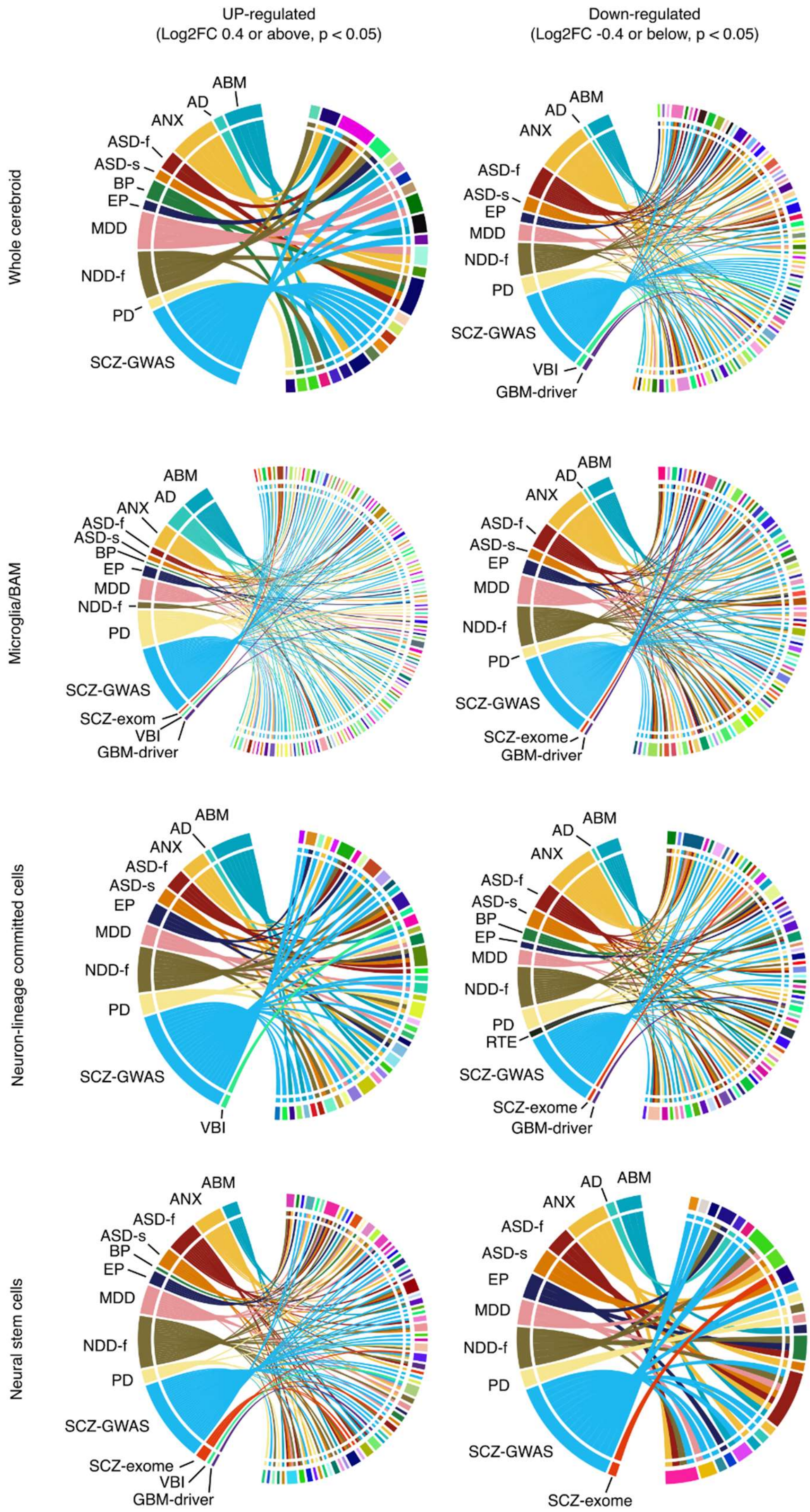
UP-regulated
(Log2FC 0.4 or above, p < 0.05)
Down-regulated
(Log2FC -0.4 or below, p < 0.05)
Whole cerebroid
ABM
AD
ANX
ASD-f
ASD-s
BP
EP
MDD
NDD-f
PD
SCZ-GWAS
ABM
AD
ANX
ASD-f
ASD-s
EP
MDD
NDD-f
PD
SCZ-GWAS
VBI
GBM-driver
Microglia/BAM
ABM
AD
ANX
ASD-f
ASD-s
BP
EP
MDD
NDD-f
PD
SCZ-GWAS
SCZ-exom
VBI
GBM-driver
ABM
AD
ANX
ASD-f
ASD-s
EP
MDD
NDD-f
PD
SCZ-GWAS
SCZ-exome
GBM-driver
Neuron-lineage committed cells
ABM
AD
ANX
ASD-f
ASD-s
EP
MDD
NDD-f
PD
SCZ-GWAS
VBI
ABM
AD
ANX
ASD-f
ASD-s
BP
EP
MDD
NDD-f
PD
RTE
SCZ-GWAS
SCZ-exome
GBM-driver
Neural stem cells
ABM
ANX
ASD-f
ASD-s
BP
EP
MDD
NDD-f
PD
SCZ-GWAS
SCZ-exome
VBI
GBM-driver
AD
ABM
ANX
ASD-f
ASD-s
EP
MDD
NDD-f
PD
SCZ-GWAS
SCZ-exome

**Fig. S12. Differentially expressed genes in IL-17A-treated cerebroids are linked to neurological disorders.**
Chord plots showing the association of a subset of differentially expressed genes (DEGs) from IL-17A-treated cerebroids (whole cerebroids and subpopulations) with various neurological disorders. The list of genes is taken from a recent study (*38*). Disorders are listed on the left, with DEGs on the right. Abbreviations: ABM (abnormal brain morphology), AD (Alzheimer's disease), ANX (anxiety disorder), ASD-f (Autism spectrum disorders, genes selected from Fu et al), ASD-s (Autism spectrum disorders, genes selected from Satterstrom et al), BP (bipolar disorder), EP (epilepsy), MDD (major depressive disorder), NDD-f (neurodevelopmental disorders, genes selected from Fu et al), PD (Parkinson's disease), RTE (response to trauma exposure), SCZ-GWAS (schizophrenia genes identified through genome-wide association studies), SCZ-exom (schizophrenia genes identified through exome sequencing), VBI (vascular brain injury), GBM-driver (glioblastoma multiforme driver genes).

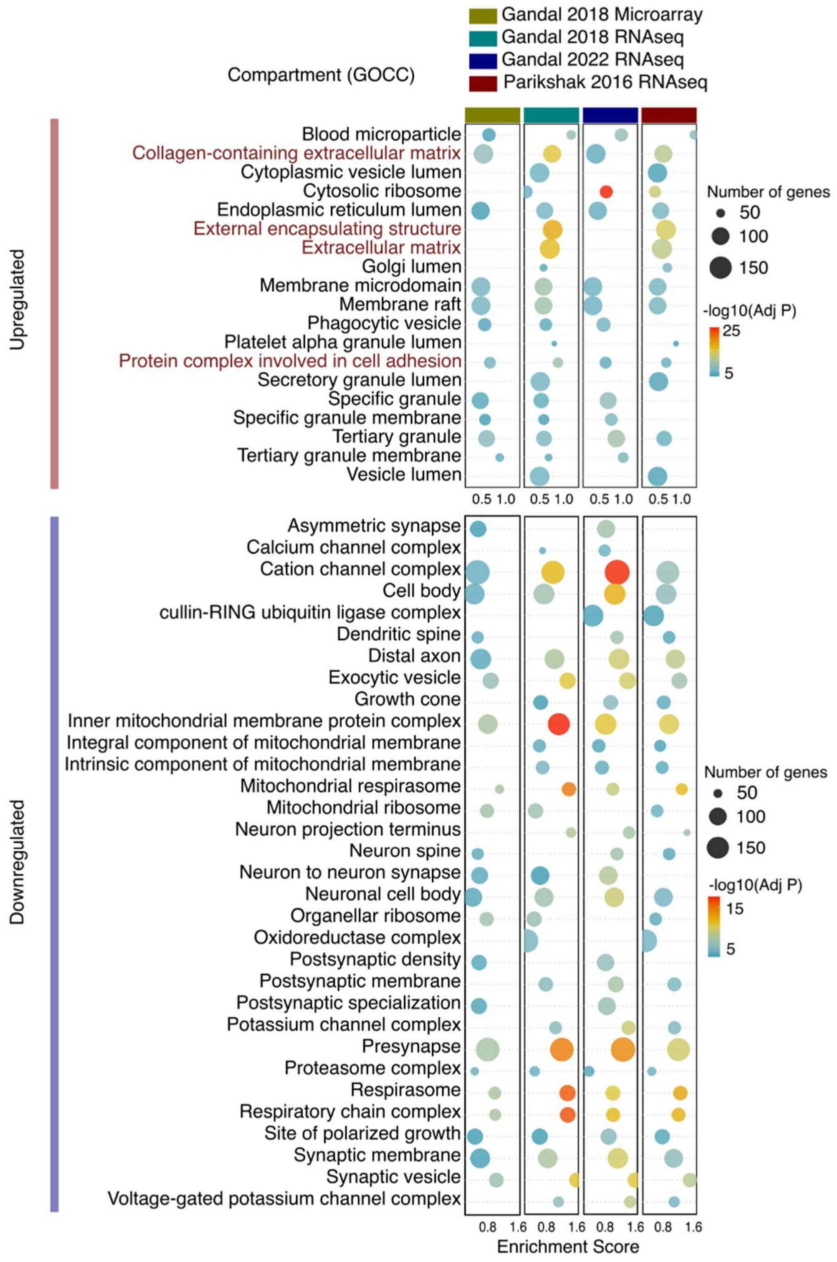


**Fig. S13. Meta-analysis of bulk transcriptomic data from post-mortem brains of individuals with ASD.**

Bubble plot displaying gene ontology cellular compartment (GOCC) enrichment for differentially regulated genes from bulk transcriptomic meta-analyses of the frontal and whole cortex of individuals with autism spectrum disorder (ASD). The analysis includes data from a meta-analysis of microarray studies and three RNA-seq studies. Enrichment analysis was performed using the STRING database, with genes ranked by Log2FC for each dataset separately. The X-axis represents the enrichment score for each term, as calculated by STRING, while columns represent individual transcriptomic datasets. Bubble size corresponds to the number of genes associated with each term, and color reflects the -log10 of the FDR-adjusted p-value. Only terms shared across two or more datasets, involving ≥50 genes, are shown. Enriched terms for both upregulated and downregulated genes are shown. ECM related terms are highlighted in red.

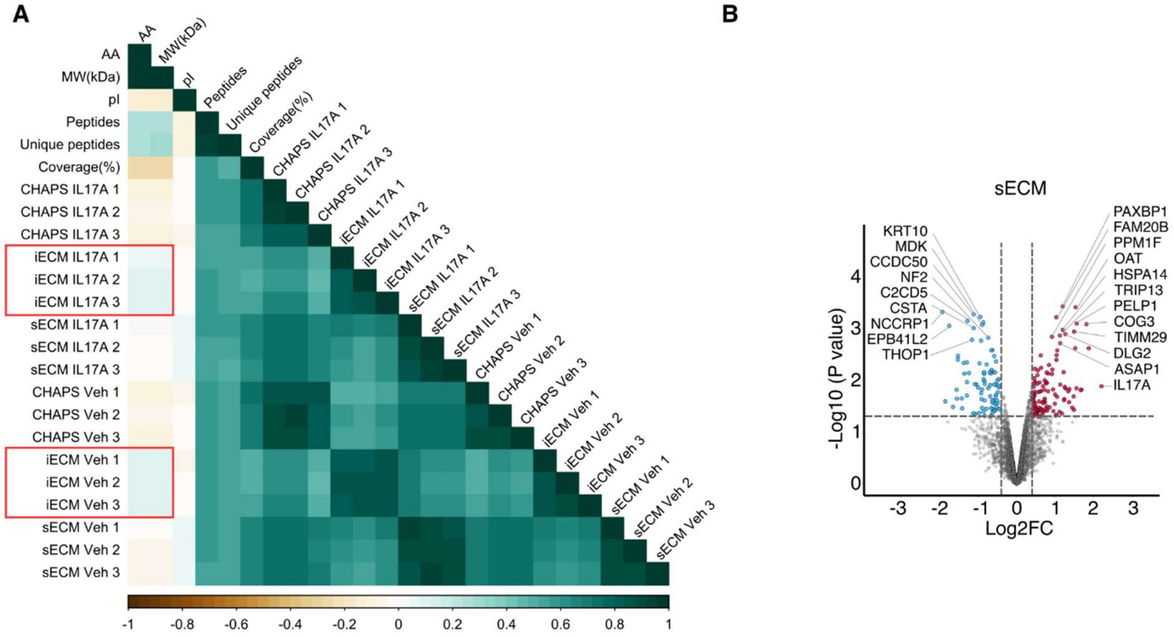


**Fig. S14. Proteomic analysis of IL-17A-treated cerebroids.**

**(A)** Correlogram showing the relationships between protein properties and their abundance across different fractions (insoluble ECM, soluble ECM, and CHAPS) from day 8 vehicle- and IL-17A-treated cerebroids. Colors indicate the strength and direction of correlations. In the insoluble ECM (iECM) fraction, protein abundance positively correlates with protein complexity, as represented by the number of amino acids (AA) and molecular weight (MW, kDa). Conversely, negative correlations are observed between protein complexity (AA and MW) and the soluble ECM (sECM) and cellular (CHAPS) fractions, suggesting enrichment of larger, more complex proteins in the iECM fraction. pI: isoelectric point.

**(B)** Volcano plot depicting differentially abundant proteins in the soluble ECM (sECM) fraction between IL-17A- and vehicle-treated day 8 cerebroids. Proteins upregulated by IL-17A treatment are shown in red, and downregulated proteins are shown in blue.

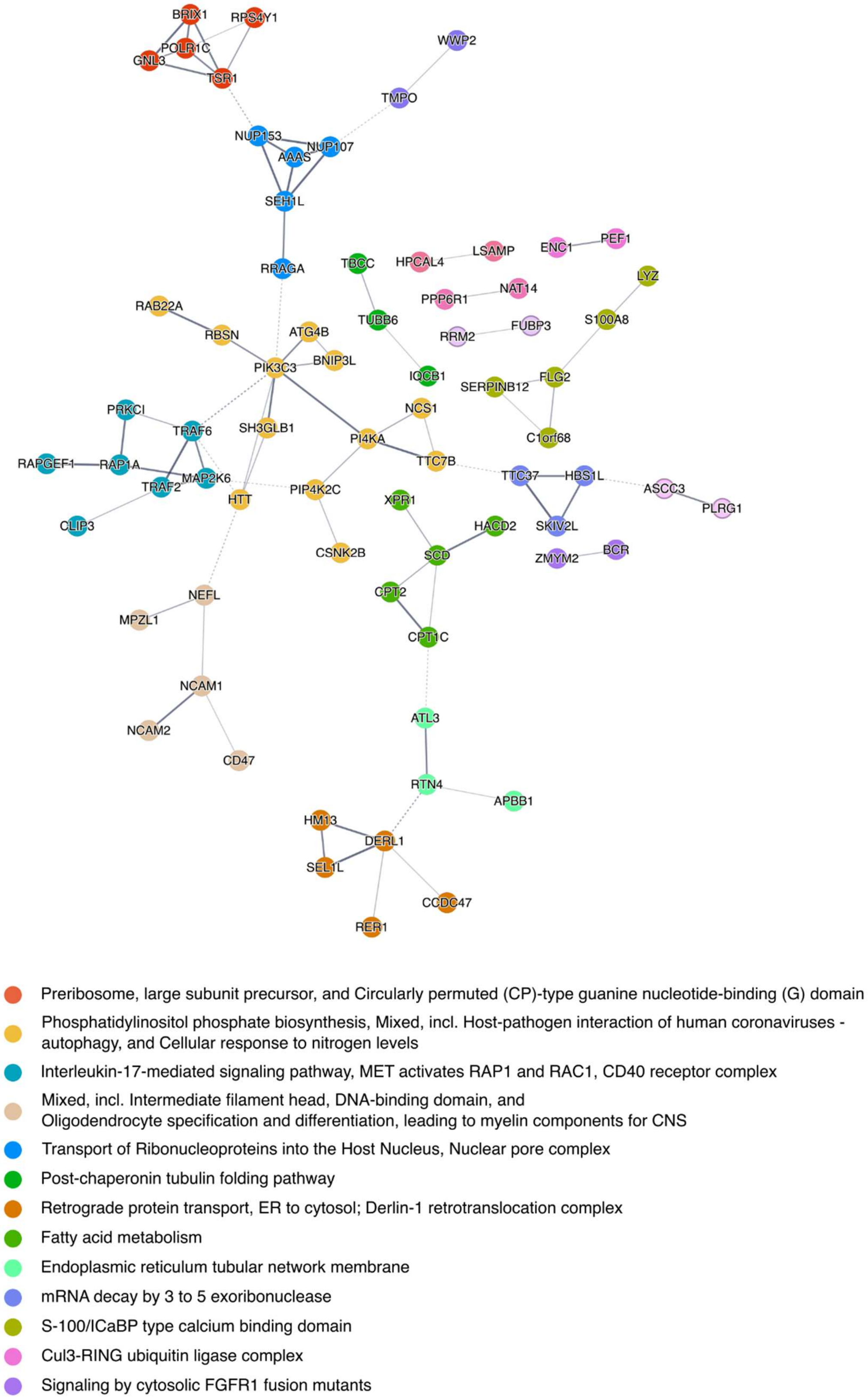
BRIX1
RPS4Y1
POLR1C
GNL3
TSR1
WWP2
TMPO
NUP153
NUP107
AAAS
SEH1L
RRAGA
TBCC
HPCAL4
LSAMP
ENC1
PEF1
LYZ
PPP6R1
NAT14
RAB22A
TUBB6
FUBP3
S100A8
RRM2
RBSN
ATG4B
BNIP3L
PIK3C3
IQCB1
SERPINB12
FLG2
PRKCI
NCS1
SH3GLB1
TRAF6
PI4KA
C1orf68
RAPGEF1
RAP1A
TTC7B
MAP2K6
TTC37
HBS1L
TRAF2
ASCC3
HTT
PIP4K2C
PLRG1
XPR1
CLIP3
HACD2
SKIV2L
BCR
CSNK2B
SCD
ZMYM2
NEFL
CPT2
MPZL1
CPT1C
NCAM1
ATL3
NCAM2
CD47
RTN4
APBB1
HM13
DERL1
SEL1L
CCDC47
RER1
Preribosome, large subunit precursor, and Circularly permuted (CP)-type guanine nucleotide-binding (G) domain
Phosphatidylinositol phosphate biosynthesis, Mixed, incl. Host-pathogen interaction of human coronaviruses - autophagy, and Cellular response to nitrogen levels
Interleukin-17-mediated signaling pathway, MET activates RAP1 and RAC1, CD40 receptor complex
Mixed, incl. Intermediate filament head, DNA-binding domain, and Oligodendrocyte specification and differentiation, leading to myelin components for CNS
Transport of Ribonucleoproteins into the Host Nucleus, Nuclear pore complex
Post-chaperonin tubulin folding pathway
Retrograde protein transport, ER to cytosol; Derlin-1 retrotranslocation complex
Fatty acid metabolism
Endoplasmic reticulum tubular network membrane
mRNA decay by 3 to 5 exoribonuclease
S-100/ICaBP type calcium binding domain
Cul3-RING ubiquitin ligase complex
Signaling by cytosolic FGFR1 fusion mutants

**Fig. S15. Protein-protein interaction network of differentially abundant proteins in the CHAPS fraction of IL-17A-treated cerebroids.**

Protein-protein interaction network of differentially abundant proteins from the CHAPS fraction of IL-17A-treated versus vehicle-treated day 8 cerebroids. The CHAPS fraction includes proteins from cellular components. The network was constructed using the STRING database, where each node represents a protein, and each line indicates a protein-protein interaction. The thickness of the lines corresponds to the confidence level of the interaction evidence. Unconnected nodes are omitted. K-means clustering was applied to the network, with each color representing a distinct protein cluster associated with an enriched biological term

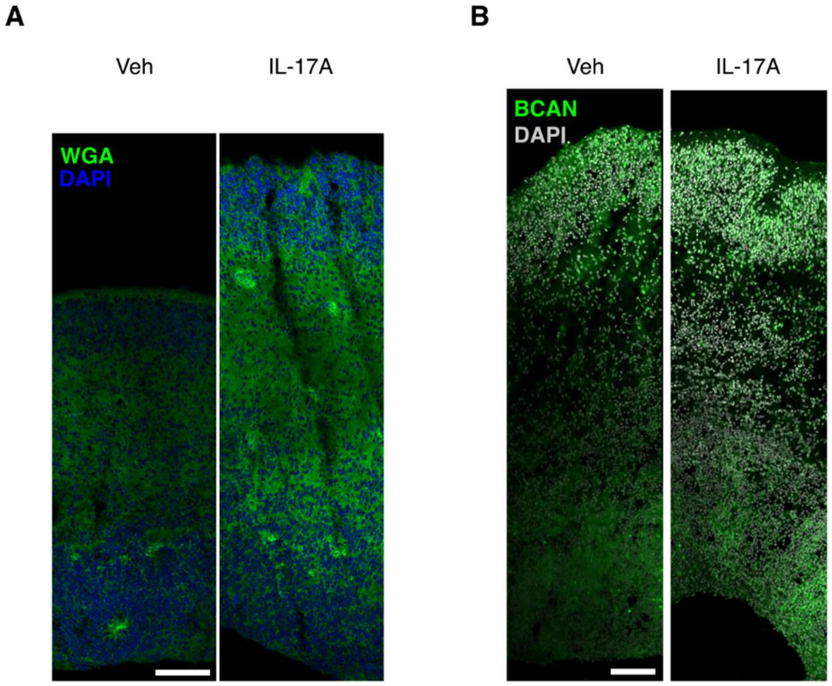


**Fig. S16. Remodeling of the extracellular matrix in IL-17A-treated cerebroids.**
**(A)** Representative fluorescence images of Day 8 cerebroids stained with Alexa fluor conjugated wheat germ agglutinin (WGA) and DAPI. WGA binds to N-acetylglucosamine and N-acetylneuraminic acid (sialic acid) residues and labels glycocalyx and glycoprotein containing ECM.
**(B)** Representative fluorescence images of Day 8 cerebroids stained for the extracellular matrix components brevican (BCAN).
Scale bar, 100 µm.

**Table S1. Donor information**

| Donor ID | Delivery to dissection time (minutes) | Age of gestation (GW+D) | Sex |
|---|---|---|---|
| 1 | 26 | 14+4 | male |
| 2 | 70 | 12+6 | male |
| 3 | 235 | 13+2 | male |
| 4 | 30 | 14+1 | male |
| 5 | 18 | 14+2 | female |
| 6 | 41 | 12 | male |
| 7 | 27 | 14+3 | male |
| 8 | 93 | 11+2 | female |
| 9 | 33 | 12+3 | male |
| 10 | 31 | 12+2 | female |
| 11 | 268 | 12+3 | male |
| 12 | 75 | 13+6 | male |
| 13 | 32 | 13+5 | male |
| 14 | 10 | 14+2 | male |
| 15 | 25 | 13+0 | male |
| 16 | 20 | 11+0 | female |

**Table S2. Primary antibodies**

| List of Primary antibodies | | | | |
|---|---|---|---|---|
| **Primary Antibody** | **Source/Isotype** | **Catalogue #** | **Dilution** | **External validation (Number of publications)** |
| **BM28/MCM2** | Mouse IgG | BD Biosciences - #610701 | 500 | 39 |
| **CD68** | Mouse IgG | Sigma - #AMAB90874 | 500 | |
| **Cleaved Caspase-3** | Rabbit IgG | Cell Signalling Technology - #9661 | 400 | 10478 |
| **Ctip2** | Rat IgG2a | Abcam - #Ab18465 | 1000 | 841 |
| **EOMES/Tbr2** | Sheep IgG | R&D Systems #AF6166 | 400 | 25 |
| **GFAP** | Mouse IgG | Millipore - #MAB360-25UL | 1000 | 65 |
| **GFAP** | Chicken IgY | Merck - AB5541 | 1000 | 299 |
| **GFP** | Goat IgG | Rockland - #600-101-215 | 1000 | 419 |
| **HOPX** | Rabbit IgG | Proteintech - #11419-1-AP | 100 | 36 |
| **HOPX** | Mouse IgG1 | Santa Cruz - #sc-398703 | 100 | 81 |
| **Iba1** | Rabbit IgG | Wako - 019-19741 | 1000 | 4528 |
| **Ki67** | Rabbit IgG | Abcam - #Ab15580 | 500 | 4218 |
| **Nestin** | Rabbit IgG | Abcam - #Ab27952 | 500 | 20 |
| **NeuN** | Mouse IgG1 | Merck - #MAB377 | 1000 | 6946 |
| **Olig-2** | Rabbit IgG | Merck - AB9610 | 500 | 1170 |
| **PTPRZ1** | Rabbit IgG | Sigma - #HPA015103 | 500 | 6 |
| **Reelin** | Mouse IgG1 | Merck - #MAB5366 | 500 | 68 |
| **S100β** | Mouse IgG1 | Merck - S2532 | 500 | 386 |
| **SATB2** | Rabbit IgG | Abcam - #ab34735 | 1000 | 90 |
| **Sox2** | Rat IgG | Invitrogen - #14-9811-82 | 1000 | 67 |
| **Sox2** | Mouse IgG1 | Abcam #Ab79351 | 1000 | 101 |
| **MAP2** | Chicken IgY | Abcam - #ab5392 | 1000 | 1135 |
| **CD31** | Rat IgG2a, κ | BioLegend - #102513 | 200 | 26 |
| **Anti-Heparan Sulfate Proteoglycan, clone A7L6** | Rat IgG2ak | Sigma-Aldrich - #MAB1948P | 200 | 46 |
| **BCAN** | Rabbit IgG | Proteintech - #19017-1-AP | 200 | 2 |
| **Anti-Chondroitin Sulfate antibody [CS-56]** | Mouse IgM | Abcam - #ab11570 | 200 | 72 |
| **COL3A** | Rabbit IgG | Proteintech - 22734-1-AP | 200 | 381 |
| **COLIVa1** | Rabbit IgG | NOVUS - NB120-6586 | 200 | 39 |
| **PDGFRB** | Rabbit IgG | Invitrogen MA5-15143 | 300 | 11 |

**Table S3. Secondary antibodies**

| List of secondary antibodies | | | | | | |
|---|---|---|---|---|---|---|
| **Species Target** | **Fluorophore** | **Ex/Em Max** | **Source/Isotype** | **Ab Format** | **Dilution** | **Catalogue #** |
| Mouse | Alexa Fluor 555 | 553/568 nm | Donkey ms-IgG (H+L) | Whole IgG | 300 | Invitrogen - A-31570 |
| Rat | Alexa Fluor 488 | 499/520 nm | Goat rat-IgG (H+L) | Whole IgG | 300 | Invitrogen - A-11006 |
| Rat | Alexa Fluor 594 | 590/618 nm | Goat rat-IgG (H+L) | Whole IgG | 300 | Invitrogen - A-11007 |
| Rat | Alexa Fluor 647 | 650/671 nm | Goat rat-IgG (H+L) | Whole IgG | 300 | Invitrogen - A-21247 |
| Sheep | Alexa Fluor 488 | 499/520 nm | Donkey sh-IgG (H+L) | Whole IgG | 300 | Invitrogen - A-11015 |
| Mouse | Alexa Fluor 488 | 499/520 nm | Goat mouse-IgG (H+L) | Whole IgG | 300 | Invitrogen - 10727464 |
| Mouse | Alexa Fluor 647 | 650/671 nm | Goat mouse-IgG (H+L) | Whole IgG | 300 | Invitrogen - 15627898 |
| Rabbit | Alexa Fluor 488 | 499/520 nm | Goat rabbit-IgG (H+L) | Whole IgG | 300 | Invitrogen - 10236882 |
| Rabbit | Alexa Fluor 594 | 590/618 nm | Goat rabbit-IgG (H+L) | Whole IgG | 300 | Invitrogen - 10266352 |
| Rabbit | Alexa Fluor 647 | 650/671 nm | Goat rabbit-IgG (H+L) | Whole IgG | 300 | Invitrogen - 15615485 |
| Mouse | Alexa Fluor 594 | 590/618 nm | Goat mouse-IgG (H+L) | Whole IgG | 300 | Invitrogen - 10524773 |
| Chicken | Alexa Fluor 488 | 499/520 nm | Goat chicken-IgY (H+L) | Whole IgG | 300 | Invitrogen - 10286672 |

**Table S4. Primers sequences**

| Gene | Forward (5'-3') | Reverse (5'-3') |
|---|---|---|
| **Sexing** | | |
| ZFX | GCACTTCTTTGGTATCTGAGAAAGT | ATAATCACATGGAGAGCCACAAGCT |
| SRY | CCCATGAACGCATTCATTGTGTGG | ATTTTAGCCTTCCGACGAGGTCGATA |
| Housekeeping | | |
| PMM1 | AACATCTCGCCCATCGGCC | TCAAAGCTGATCATGCCTCCTCG |
| RPLP0 | TCTACAACCCTGAAGTGCTTGAT | CAATCTGCAGACAGACACTGG |
| SDHA | ACGTCACGAAGGAGCCGATCC | ATGTACCGAGGCACAGGCGG |
| **Markers of inflammation** | | |
| IL1B | ATGCACCTGTACGATCACTG | ACAAAGGACATGGAGAACACC |
| IL6 | CCACTCACCTCTTCAGAACG | CATCTTTGGAAGGTTCAGGTTG |
| TNF | GTCAACCTCCTCTCTGCCAT | CCAAAGTAGACCTGCCCAGA |
| NOS2 | GCCCTCACCTACTTCCTG | ACTTCCACTTGCTGTACTCTG |
| C3 | AGTCTTTGTACGTGTCTGCC | ACTTGGGTGTCTTGGTGAAG |

### References


1. E. Bongaerts *et al.*, Maternal exposure to ambient black carbon particles and their presence in maternal and fetal circulation and organs: an analysis of two independent population-based observational studies. *The Lancet Planetary Health* **6**, e804-e811 (2022).
2. P. J. O'Shaughnessy *et al.*, Developmental Changes in Human Fetal Testicular Cell Numbers and Messenger Ribonucleic Acid Levels during the Second Trimester. *The Journal of Clinical Endocrinology & Metabolism* **92**, 4792-4801 (2007).
3. A. McQuade *et al.*, Development and validation of a simplified method to generate human microglia from pluripotent stem cells. *Mol Neurodegener* **13**, 67 (2018).
4. F. Ye *et al.*, DISC1 Regulates Neurogenesis via Modulating Kinetochore Attachment of Ndel1/Nde1 during Mitosis. *Neuron* **96**, 1041-1054 e1045 (2017).
5. X. Qian *et al.*, Brain-Region-Specific Organoids Using Mini-bioreactors for Modeling ZIKV Exposure. *Cell* **165**, 1238-1254 (2016).
6. K. Alves de Lima *et al.*, Meningeal gammadelta T cells regulate anxiety-like behavior via IL-17a signaling in neurons. *Nat Immunol* **21**, 1421-1429 (2020).
7. A. M. McGinley *et al.*, Interleukin-17A Serves a Priming Role in Autoimmunity by Recruiting IL-1beta-Producing Myeloid Cells that Promote Pathogenic T Cells. *Immunity* **52**, 342-356 e346 (2020).
8. P. Konieczny *et al.*, Interleukin-17 governs hypoxic adaptation of injured epithelium. *Science* **377**, eabg9302 (2022).
9. M. D. Reed *et al.*, IL-17a promotes sociability in mouse models of neurodevelopmental disorders. *Nature* **577**, 249-253 (2020).
10. L. Sun *et al.*, IL-10 Dampens an IL-17-Mediated Periodontitis-Associated Inflammatory Network. *J Immunol* **204**, 2177-2191 (2020).
11. M. Bambouskova *et al.*, Electrophilic properties of itaconate and derivatives regulate the IkappaBzeta-ATF3 inflammatory axis. *Nature* **556**, 501-504 (2018).
12. R. Bechara *et al.*, The m(6)A reader IMP2 directs autoimmune inflammation through an IL-17- and TNFalpha-dependent C/EBP transcription factor axis. *Sci Immunol* **6**, (2021).
13. P. Pavlidis *et al.*, Cytokine responsive networks in human colonic epithelial organoids unveil a molecular classification of inflammatory bowel disease. *Cell Rep* **40**, 111439 (2022).
14. D. Sinkeviciute, A. Aspberg, Y. He, A. C. Bay-Jensen, P. Onnerfjord, Characterization of the interleukin-17 effect on articular cartilage in a translational model: an explorative study. *BMC Rheumatol* **4**, 30 (2020).
15. N. Walker, P. Filis, P. J. O'Shaughnessy, M. Bellingham, P. A. Fowler, Nutrient transporter expression in both the placenta and fetal liver are affected by maternal smoking. *Placenta* **78**, 10-17 (2019).
16. E. A. Bordt *et al.*, Isolation of Microglia from Mouse or Human Tissue. *STAR Protocols* **1**, 100035-100035 (2020).
17. S. Chen, Ultrafast one-pass FASTQ data preprocessing, quality control, and deduplication using fastp. *iMeta* **2**, (2023).
18. R. Patro, G. Duggal, M. I. Love, R. A. Irizarry, C. Kingsford, Salmon provides fast and bias-aware quantification of transcript expression. *Nat Methods* **14**, 417-419 (2017).

19. M. I. Love *et al.*, Tximeta: Reference sequence checksums for provenance identification in RNA-seq. *PLoS Comput Biol* **16**, e1007664 (2020).
20. M. I. Love, W. Huber, S. Anders, Moderated estimation of fold change and dispersion for RNA-seq data with DESeq2. *Genome Biol* **15**, 550 (2014).
21. M. C. McCabe, A. J. Saviola, K. C. Hansen, Mass Spectrometry-Based Atlas of Extracellular Matrix Proteins across 25 Mouse Organs. *Journal of Proteome Research* **22**, 790-801 (2023).
22. M. E. Ritchie *et al.*, limma powers differential expression analyses for RNA-sequencing and microarray studies. *Nucleic Acids Res* **43**, e47 (2015).
23. K. Garbett *et al.*, Immune transcriptome alterations in the temporal cortex of subjects with autism. *Neurobiology of Disease* **30**, 303-311 (2008).
24. I. Voineagu *et al.*, Transcriptomic analysis of autistic brain reveals convergent molecular pathology. *Nature* **474**, 380-384 (2011).
25. M. L. Chow *et al.*, Age-Dependent Brain Gene Expression and Copy Number Anomalies in Autism Suggest Distinct Pathological Processes at Young Versus Mature Ages. *PLoS Genetics* **8**, e1002592-e1002592 (2012).
26. S. Gupta *et al.*, Transcriptome analysis reveals dysregulation of innate immune response genes and neuronal activity-dependent genes in autism. *Nature Communications* **5**, 5748-5748 (2014).
27. N. N. Parikshak *et al.*, Genome-wide changes in lncRNA, splicing, and regional gene expression patterns in autism. *Nature* **540**, 423-427 (2016).
28. C. Wright *et al.*, Altered expression of histamine signaling genes in autism spectrum disorder. *Translational Psychiatry* **7**, e1126-e1126 (2017).
29. M. J. Gandal *et al.*, Shared molecular neuropathology across major psychiatric disorders parallels polygenic overlap. *Science* **359**, 693-697 (2018).
30. M. J. Gandal *et al.*, Broad transcriptomic dysregulation occurs across the cerebral cortex in ASD. *Nature* **611**, 532-539 (2022).
31. P. Zhang *et al.*, Neuron-specific transcriptomic signatures indicate neuroinflammation and altered neuronal activity in ASD temporal cortex. *Proceedings of the National Academy of Sciences* **120**, (2023).
32. D. Velmeshev *et al.*, Single-cell genomics identifies cell type–specific molecular changes in autism. *Science* **364**, 685-689 (2019).
33. M. J. Herrero *et al.*, Identification of amygdala-expressed genes associated with autism spectrum disorder. *Molecular Autism* **11**, 39-39 (2020).
34. D. Szklarczyk *et al.*, The STRING database in 2023: protein–protein association networks and functional enrichment analyses for any sequenced genome of interest. *Nucleic Acids Research* **51**, D638-D646 (2023).
35. D. Szklarczyk *et al.*, The STRING database in 2021: customizable protein–protein networks, and functional characterization of user-uploaded gene/measurement sets. *Nucleic Acids Research* **49**, D605-D612 (2021).
36. Z. Xie *et al.*, Gene Set Knowledge Discovery with Enrichr. *Curr Protoc* **1**, e90 (2021).
37. H. Han *et al.*, TRRUST v2: an expanded reference database of human and mouse transcriptional regulatory interactions. *Nucleic Acids Res* **46**, D380-D386 (2018).
38. S. Kim *et al.*, An integrative single-cell atlas to explore the cellular and temporal specificity of neurological disorder genes during human brain development. *bioRxiv*, (2024).
